# Mechanisms of nuclear segregation in a multinucleate multibudding yeast

**DOI:** 10.1101/2025.04.14.648742

**Authors:** Claudia A. Petrucco, Alex W. Crocker, Alison C.E. Wirshing, Analeigha V. Colarusso, Maya Waarts, Amy S. Gladfelter, Daniel J Lew

**Affiliations:** Department of Cell Biology, Duke University School of Medicine, Durham, NC 27710; Biology Department, Massachusetts Institute of Technology, Cambridge, MA 02139; Department of Molecular and Cellular Biology, Harvard University, Cambridge, MA 02138

## Abstract

Budding yeasts present an especially challenging geometry for segregation of chromosomes, which must be delivered across the narrow mother-bud neck into the bud. Studies in the model yeast *Saccharomyces cerevisiae* have revealed an elaborate set of mechanisms that selectively orient one mitotic spindle pole towards the bud and then drive spindle elongation along the mother-bud axis, ensuring nuclear segregation between mother and bud. It is unclear how these pathways might be adapted to yield similar precision in more complex cell geometries. Here we provide the first description of the dynamics of mitosis in a multi-nucleate, multibudding yeast, *Aureobasidium pullulans,* and identify many unexpected differences from uninucleate yeasts. Mitotic spindles do not orient along the mother-bud axis prior to anaphase, and accurate nuclear segregation often occurs after spindle disassembly. Cortical Num1-dynein forces pull highly mobile nuclei into buds, and once a nucleus enters a bud, it discourages others from entering, ensuring that most daughters inherit only one nucleus.

## Introduction

Proliferating eukaryotic cells coordinate cytokinesis with mitosis both temporally (mitosis first, then cytokinesis) and spatially (division furrow perpendicular to spindle axis) to produce viable daughters (Glotzer, 2017). Spindle orientation is influenced by several spatial cues, including cell shape, cortical tension, adjacent cells, and various pre-localized polarity determinants. Correct orientation is critical for development and tissue patterning in plants and animals (Morin and Bellaiche, 2011; Lipka et al., 2015).

Studies of model yeasts have been instrumental in revealing the detailed mechanisms of spindle orientation (Segal and Bloom, 2001; Varshney and Sanyal, 2019). Unlike in many symmetrically dividing cells, the site of cytokinesis in budding yeasts (the mother-bud neck) is specified prior to spindle formation. Thus, spindle orientation along the pre-determined mother-bud axis is needed to ensure that both mother and bud inherit a nucleus. Delivering one pole of the mitotic spindle across a narrow mother-bud neck creates a significant challenge for both geometrical and mechanical reasons (Haase and Lew, 2007). The area of the neck aperture (about 1 μm diameter) represents only 1-2% of the mother cell’s surface, so a spindle pole would be unlikely to encounter the aperture without guidance (Kubo et al., 2019). Because most yeasts undergo a closed mitosis, even if a spindle pole does encounter the opening, some force would be required to squeeze the larger nucleus through the smaller bud neck (De Souza and Osmani, 2009). In the model budding yeast, *Saccharomyces cerevisiae,* these problems are solved through pathways that deliver astral microtubule plus ends into the bud to pre-orient the spindle and then pull the attached nucleus through the neck during anaphase.

The molecular basis for spindle orientation in *S. cerevisiae* has been extensively investigated. Early in the cell cycle, before spindle assembly, a protein called Kar9 (Miller and Rose, 1998) is delivered to cytoplasmic microtubule plus ends, where it links the plus-end microtubule-binding EB1-related protein Bim1 to the type V myosin Myo2 (Lee et al., 2000; Yin et al., 2000). Using polarized actin cables, the myosin delivers the microtubule plus ends into the bud, drawing the attached spindle pole bodies, and hence the nucleus, towards the neck (Theesfeld et al., 1999; Hwang et al., 2003). In mitosis, the microtubule minus-end motor dynein is delivered to microtubule plus ends and off-loaded to cortical domains containing the anchor protein Num1 (Lee et al., 2003; Farkasovsky and Kuntzel, 2001; Markus and Lee, 2011). Num1 is thought to play an analogous role to the animal protein NuMA (Greenberg et al., 2018).

Dynein then drags the cortical microtubules along the cortex to pull the attached spindle pole body through the neck. Forces from kinesin V motors in the spindle midzone also act to elongate the spindle, pushing one pole into the bud and the other to the back of the mother cell (Saunders et al., 1995). Together, these pathways pre-orient the mitotic spindle towards the bud and promote spindle elongation through the mother-bud neck to deliver one daughter nucleus to the bud while keeping the other in the mother cell.

The importance of correctly orienting the spindle is highlighted by the finding that *S. cerevisiae* cells appear to check that nuclear segregation has successfully delivered one nucleus to the bud before permitting mitotic exit (Yeh et al., 1995). This pathway, called the spindle position checkpoint, links the activation of a signaling pathway (the mitotic exit network) to spindle pole position (Zhou et al., 2024). When errors occur and anaphase spindles elongate entirely in the mother cell, the checkpoint maintains a cell-cycle arrest in anaphase until the spindle reorients and delivers one pole into the bud. Combined with the partitioning mechanisms discussed above, this failsafe assures that yeast do not produce anucleate buds.

The geometric and mechanical challenges discussed above are likely to apply to many or all budding yeasts. However, an additional challenge arises for budding yeasts that are multinucleate and make more than one bud during a cell cycle, like the pathogens *Paracoccidioides brasiliensis* (Peçanha et al., 2022), and *Mucor circinelloides* (Lübbehüsen *et al*., 2003). An example of this strategy is provided by the polyextremotolerant black yeast, *Aureobasidium pullulans* (Gostincar et al., 2014). *A. pullulans* mother cells contain anywhere from one to over 30 nuclei, and they make quite variable numbers of buds (Mitchison-Field et al., 2019; Goshima, 2022; Petrucco et al., 2024; Wirshing et al., 2024). The nuclei undergo a synchronous, semi-open mitosis that seems to deliver nuclei to most if not all buds (Petrucco et al., 2024). This mode of proliferation raises the question of how the nuclei come to be appropriately segregated.

If the mechanisms studied in *S. cerevisiae* were at play in *A. pullulans*, those pathways could deliver nuclei to buds, but it is unclear how the nuclei would be apportioned among sister buds. For example, one might expect to see a subset of buds get multiple nuclei while others get none, which would seem unproductive. Thus, examination of this question in *A. pullulans* could reveal additional partitioning mechanisms, beyond those understood from *S. cerevisiae*. Alternatively, one could imagine that instead of specific mechanisms to orient mitotic spindles towards each bud, *A. pullulans* might simply distribute their nuclei evenly within the available space, as seen in some other multinucleated cells (Padilla et al., 2022). For example, nuclei are often distributed evenly within multinucleate hyphae, which seems to involve microtubule-based interactions between neighboring nuclei (Morris, 2003; Anderson et al., 2013; Xiang, 2019).

Such a strategy might conceivably enable nuclear segregation in multibudded cells, without specific spindle orientation pathways, although the multibudded morphology, with several narrow necks separating the buds from the mother, presents more challenges for nuclear partitioning than the tubular hyphal morphology.

Here we describe microtubule behavior and analyze nuclear segregation mechanisms in *A. pullulans*. Our findings suggest that nuclear segregation in these cells requires neither spindle pre-alignment nor any obvious coordination between spindles. Instead, nuclei are delivered into buds by Num1-dependent pulling of spindle pole bodies, and once one nucleus enters a bud, it hinders the entry of other nuclei. This results in a relatively even distribution of nuclei between mother and buds where each bud receives at least, and usually only, one nucleus.

## Results

### Fidelity of nuclear inheritance in *A. pullulans*

To visualize nuclear dynamics in *A. pullulans*, we imaged proliferating cells expressing a fluorescently-tagged histone (Petrucco et al., 2024). Consistent with previous findings on fixed cells (Mitchison-Fields et al., 2019), almost all multinucleate mothers made either the same number of buds as they had nuclei (48%, n = 587 multinucleate cells), or they made fewer buds than nuclei (51%, n = 587 multinucleate cells). There were, however, rare cases in which mothers made more buds than they had nuclei (Fig. 1A,B). During mitosis, sister buds from the same mother usually received the same number of nuclei, though there were exceptions (Fig. 1C,D). A majority of buds received a single nucleus, but a substantial minority received two nuclei. On rare occasions a bud received >2 nuclei, and it was very rare for buds to receive no nuclei (Fig. 1E,F). Interestingly, the buds that received two nuclei were on average larger than those that received only one (Fig. 1G). The rare anucleate buds arose either from instances where mother cells had made too many buds for the available nuclei (4 of 1934 buds observed) or from an apparent mitotic segregation error (1 of 1934 buds observed). Similar results were obtained with a strain expressing a fluorescent marker fused to a nuclear localization sequence (NLS) (Petrucco et al., 2024) instead of a histone (Fig. 1F). These findings indicate that *A. pullulans* has a robust nuclear segregation mechanism to ensure that each bud receives at least one nucleus, despite the complex situation of many nuclei with many potential destination buds.

**Figure 1:**
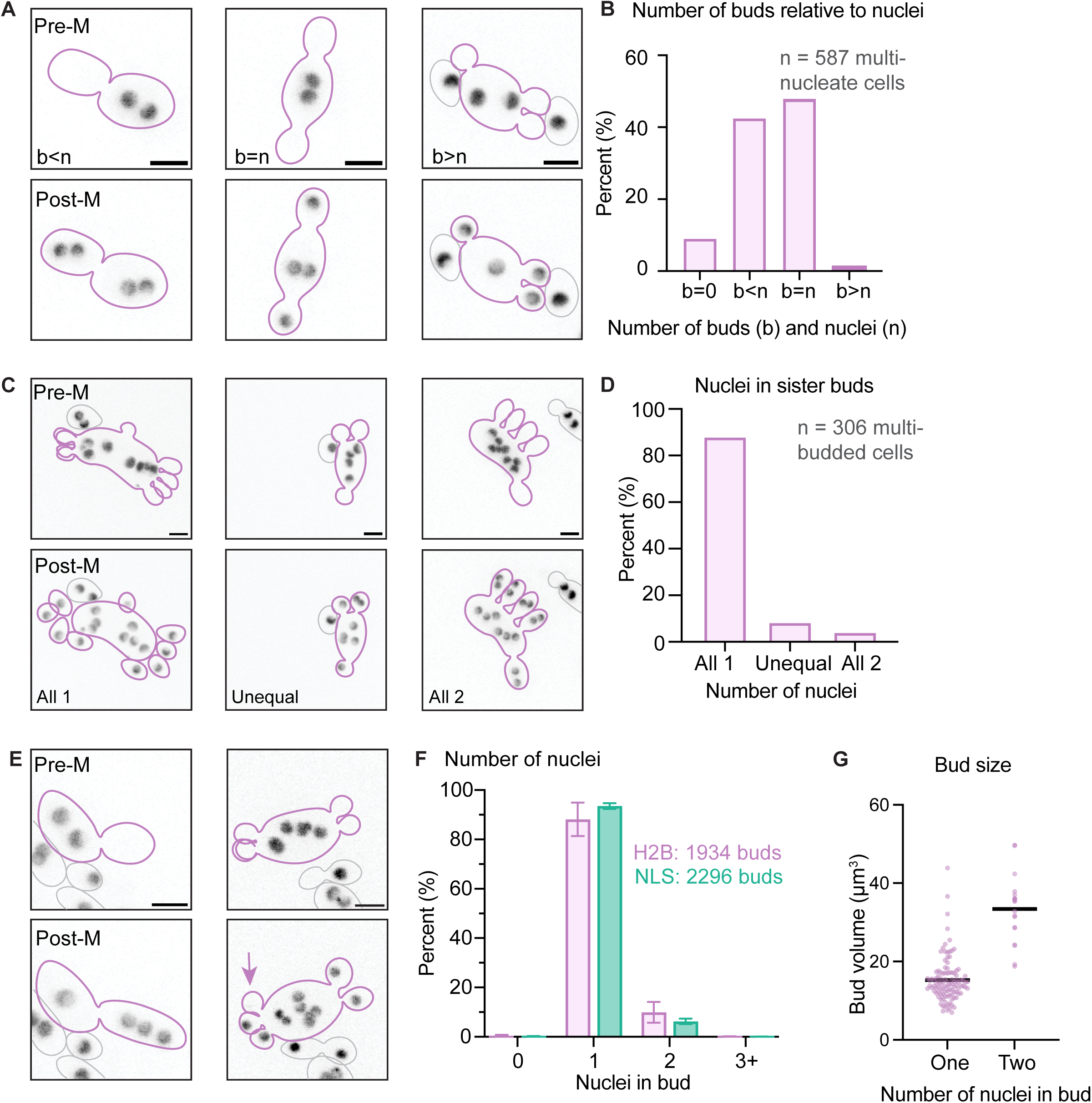
Nuclear inheritance. (A) Inverted maximum projection images of binucleate mother cells making different numbers of buds. Upper and lower panels show the same highlighted cell before and after mitosis. (B) Number of buds produced (b) in relation to the number of pre-mitotic nuclei (n) in the mother cell (n= 587 multinucleate cells with 2 or more nuclei, as visualized with a histone probe). (C) Examples (as in A) of nuclear inheritance by sister buds made in the same cell cycle. (D) Number of nuclei inherited by sister buds (n= 306 mothers making two or more buds). (E) Examples (as in A) of rare cases where buds received three (left) or zero (right, arrow) nuclei. (F) Number of nuclei inherited by each bud following mitosis (pink: histone probes, n=1934 buds; green: NLS-tdTomato probe, n=2296 buds). (G) Buds that receive two nuclei are larger on average than those that receive a single nucleus. Dot plots of bud volume. Black bar indicates mean (n = 126 buds). Images and quantification from time-lapse movies of DLY23973 and DLY24568. Scale bars, 5 μm.

### Microtubules and the mitotic spindle in *A. pullulans*

We next examined how spindles and astral microtubules behave through the cell cycle. We previously showed that *A. pullulans* undergoes a semi-open mitosis in which an apparently intact nuclear envelope encases each nucleus, but nuclear pores become permeable (Petrucco et al., 2024). Similar semi-open mitoses were first described in the related *Aspergillus nidulans* (De Souza et al., 2004). This suggests that each nucleus must develop its own intranuclear spindle, and that somehow the spindles must be spatially coordinated to deliver daughter nuclei to each and every bud. To visualize microtubules in live cells, we N-terminally tagged *T*ub2 (β-tubulin) with three copies of a codon-optimized GFP. Tandem GFPs allowed brighter labeling without detectably impairing colony growth (Fig. S1).

Both interphase and mitotic networks of microtubules were detectable. A robust interphase network extended throughout the cytoplasm of the mother cell, with microtubules reaching near the tips of each bud (Fig. 2A, Fig. S2). In mitosis, cells exhibited either short or long spindles, with one bar-shaped spindle in each nucleus (Fig. 2B,C, Fig. S2, Supplemental Video 1). Longer spindles had astral microtubules extending from either pole. Other cells, which we show below were post-anaphase, displayed long but fainter microtubules emanating from discrete foci, without a spindle (Fig. 2D, Fig. S2). Spindles of different lengths could be assigned to sequential cell cycle stages (metaphase, spindle < 3 μm; anaphase, spindle > 3 μm)(Fig. 2) based on detecting the metaphase-anaphase transition by time-lapse microscopy (Supplemental Video 1). To ascertain the polarity of the microtubules, we fluorescently tagged the endogenous *A. pullulans* microtubule plus-tip protein Bim1 (EB1-related) (Berrueta et al., 1998). Bim1 puncta were visible at the tips of growing interphase microtubules (Fig. 2E), and at the tips of astral microtubules extending away from the spindle poles (Fig. 2F)(Supplemental Video 2). In summary, *A. pullulans* cells remodel an elaborate interphase cytoplasmic microtubule network to generate intranuclear spindles and extend astral cytoplasmic microtubules towards the cell periphery.

**Figure 2:**
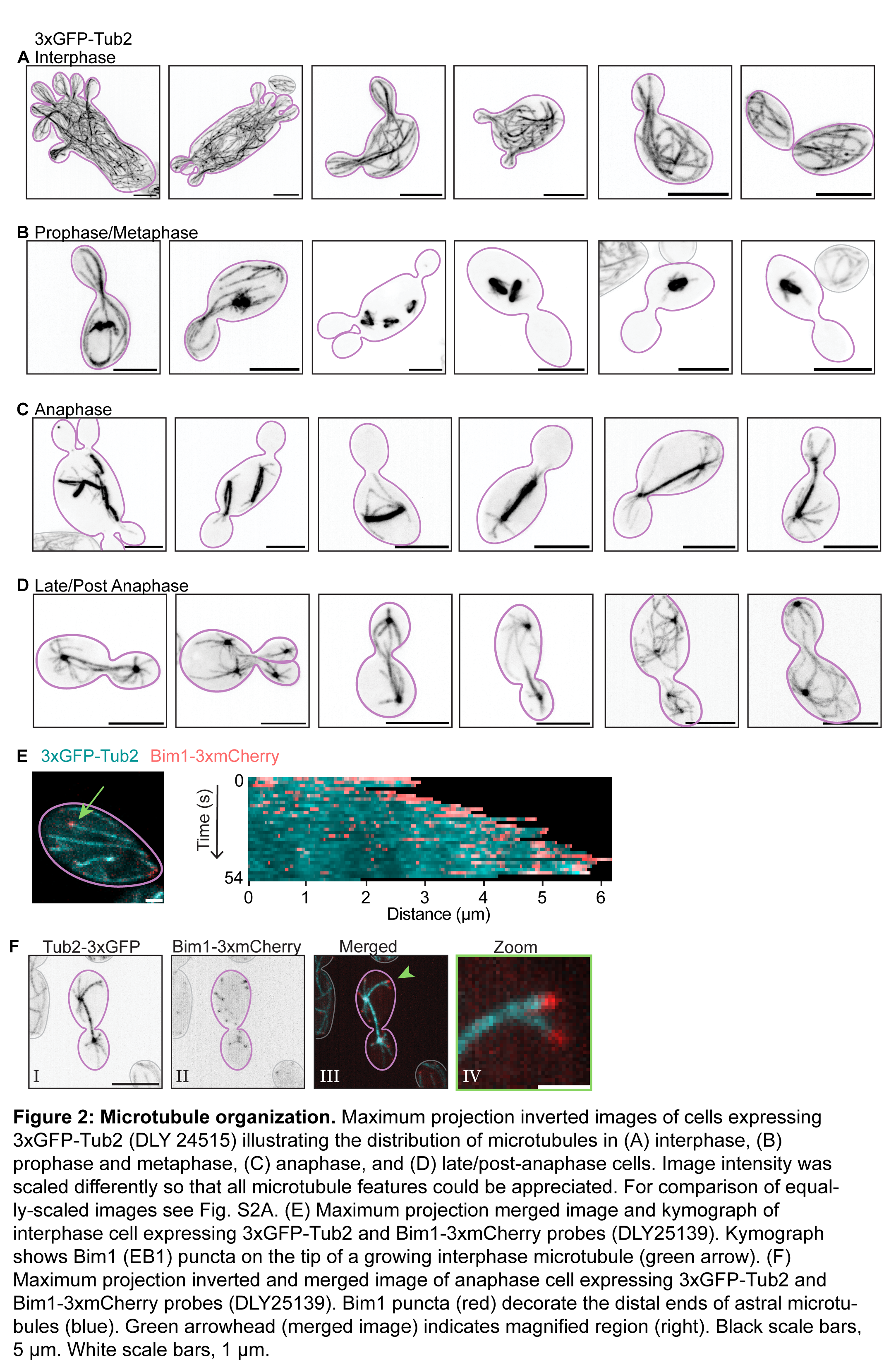
**Microtubule organization**. Maximum projection inverted images of cells expressing 3xGFP- Tub2 (DLY 24515) illustrating the distribution of microtubules in (A) interphase, (B) prophase and metaphase, (C) anaphase, and (D) late/post-anaphase cells. Image intensity was scaled differently so that all microtubule features could be appreciated. For comparison of equally-scaled images see Fig. S2A. (E) Maximum projection merged image and kymograph of interphase cell expressing 3xGFP-Tub2 and Bim1-3xmCherry probes (DLY25139). Kymograph shows Bim1 (EB1) puncta on the tip of a growing interphase microtubule (green arrow). (F) Maximum projection inverted and merged image of anaphase cell expressing 3xGFP-Tub2 and Bim1-3xmCherry probes (DLY25139). Bim1 puncta (red) decorate the distal ends of astral microtubules (blue). Green arrowhead (merged image) indicates magnified region (right). Black scale bars, 5 μm. White scale bars, 1 μm.

The interphase microtubule network was rapidly disassembled concomitant with spindle formation in a span of 1.3 ± 0.5 min (mean ± SD, n = 84 cells) (Fig. 2B, 3A)(Supplemental Video 3). Short metaphase spindles were present for 4.1 ± 1.1 min (n = 84 cells) prior to spindle elongation. This is much shorter than metaphase in *S. cerevisiae* (about 16 min) (Shaw et al., 1998) but comparable to *A. nidulans* where mitosis is completed within 5 min (Bergen and Morris, 1983; Szewczyk and Oakley, 2011). Metaphase spindles had very few astral microtubules (1.3 ± 1.6 astral microtubules, n = 62 metaphase spindles) which were short in length (1.1 ± 0.6 μm, n = 82 astral microtubules) (Fig 3B,C). As cells progressed into anaphase, the number (6.7 ± 2.3 astral microtubules, n = 47 anaphase spindles) and length (1.8 ± 1.1 μm, n = 313 astral microtubules) of astral microtubules increased. Each spindle pole was decorated with astral microtubules of varying lengths (Fig. 3B,C). Spindles became thinner as they lengthened, and it was often difficult to discern the exact time of spindle disassembly (Supplemental Video 4). The period from the onset of spindle elongation to the time when the spindle reached 6 μm (in many cases the longest structure we could confidently identify as an intact spindle) was 1.5 ± 1.6 min (n = 84 cells)(Fig. 3A). Again, this duration is much shorter than anaphase in *S. cerevisiae* (16 – 30 min) (Yeh et al., 1995; Yang et al., 1997), but consistent with the short mitosis in *A. nidulans* (5 min) (Bergen and Morris, 1983; Szewczyk and Oakley, 2011).

**Figure 3:**
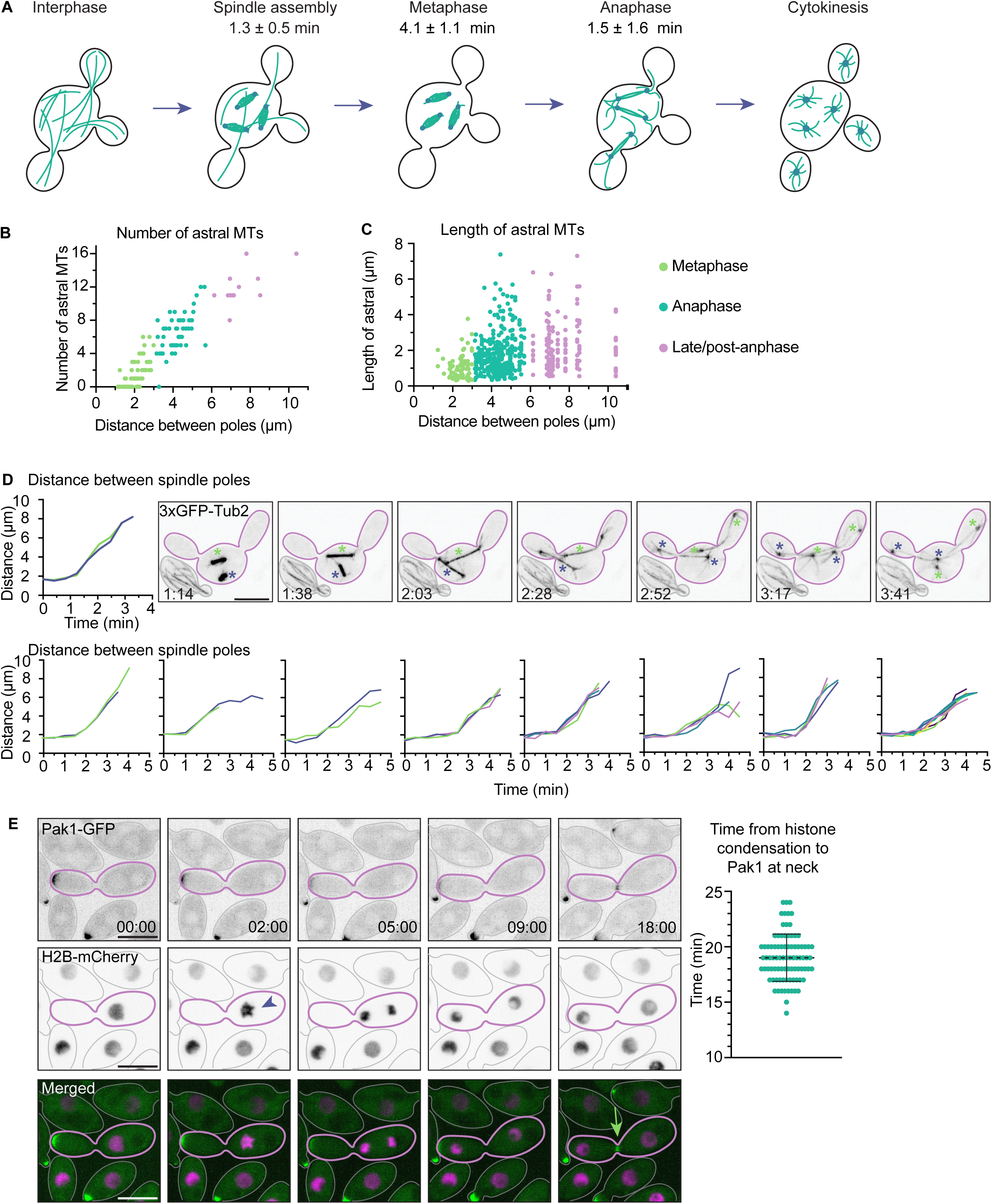
Mitotic spindle and astral microtubule dynamics. (A) Timing of mitotic events, determined from time-lapse movies of cells expressing 3xGFP-Tub2 (DLY24515). Spindle assembly was coincident with the disassembly of the interphase network of microtubules, and short (∼3 μm) bright metaphase spindles lengthened and became thinner and fainter during anaphase. Measurements from 84 cells. (B,C) The number and length of astral microtubules. Metaphase spindles had very few astral microtubules, and those were less than 3 μm long. As the spindle elongated, the number (B) and length (C) of astral microtubules increased. This continued in late/post anaphase. Measurements from 121 spindles and 539 astral microtubules. (D) Time-lapse inverted maximum projection images of cells expressing 3xGFP-Tub2 (DLY 24515). Spindles in the same cell elongated simultaneously. Each spindle and associated poles are denoted by a green or purple asterisk. Graphs plot pole-to-pole distance for each color-coded spindle during mitosis in multi-nucleate cells. (E) Timing from the onset of mitosis (chromosome condensation, purple arrowhead) to the onset of cytokinesis (Pak1 relocation to neck, green arrow) was determined from time-lapse movies of cells expressing H2B-mCherry and Pak1- GFP (CPY519). Graph: each dot indicates one cell. Mean timing is indicated with dashed line. Scale bars, 5 μm. Time is in minutes: seconds.

In multi-nucleate *A. pullulans* cells, all nuclei undergo synchronous (within 30s) chromosome condensation and decondensation (Petrucco et al., 2024). Consistently, spindles in multi-nucleate cells elongated simultaneously (Fig. 3D). Spindle poles remained highly mobile after spindle disassembly (Supplemental Video 4), and the number (12 ± 2.3 astral microtubules, n = 12 pairs of spindle poles) and length (2.2 ± 1.3 μm, n = 144 astral microtubules) of astral microtubules continued to increase (Fig. 3B,C). We used a fluorescently labeled Pak1, a polarity protein that relocates to the bud neck at the end of the cell cycle (Crocker et al., 2025), to estimate the timing of cytokinesis. The time from the onset of mitosis (chromosome condensation) until Pak1 accumulated at the neck was 19 ± 2.1 min (n = 89 cells)(Fig. 3E). Combined with the observations above, this indicates a post-anaphase interval of about 14 min prior to cytokinesis. The spindle poles remained highly mobile throughout the interval. Taken together, these findings indicate that upon anaphase onset, astral microtubules elongate and drive the rapid motility of anaphase spindles and post-anaphase nuclei.

### Metaphase spindles do not pre-orient towards the buds

We next analyzed spindle orientation to determine whether, as in *S. cerevisiae*, the spindles orient towards buds before anaphase. Most *A. pullulans* spindles did not display detectable pre-orientation towards a specific bud (n = 72 spindles, Fig 4A,B). This is consistent with the observation that metaphase spindles do not have astral microtubules long enough to reach the bud (Fig 2B,3C). In mononucleate, single-budded *A. pullulans* cells, we could unambiguously call the direction in which a spindle “should” orient, and a slight bias became evident at about 5 s before anaphase (Fig. 4A). In multinucleate cells with multiple buds, we detected no pre-orientation of spindles. In these cells, we measured the angle of the spindle in regards to the bud it did enter and the buds it did not enter, we tracked the bud which a spindle pole eventually entered but detected no preferential pre-orientation toward that bud as opposed to other buds (Fig. 4B). Tracking spindle orientation at 5 s intervals over 2 min spanning anaphase onset, we observed very dynamic and variable behaviors. In some cells, the spindle became better oriented just prior to anaphase (Fig. 4C, top). In other cells, misoriented spindles did not change orientation, even during anaphase (Fig. 4C, bottom). And in some cells, the spindle was at some point aligned but then became misaligned prior to or during anaphase (see examples in Fig. 4D). These observations indicate that spindle pre-orientation before anaphase, if it occurs at all, is the exception rather than the rule.

**Figure 4:**
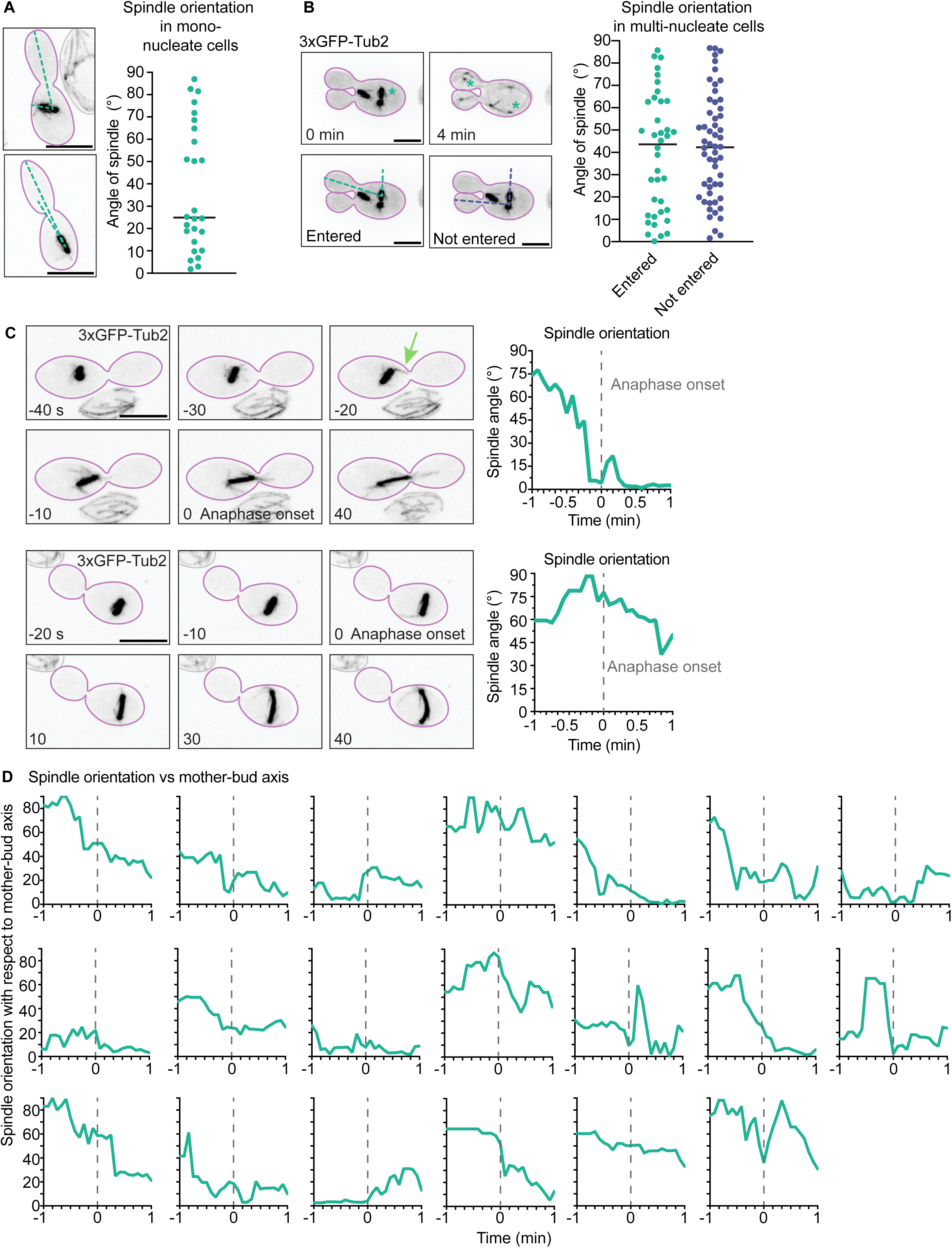
**Spindle orientation.** (A) Spindle orientation in mononucleate cells. Left: Example spindles indicating angle between spindle axis and mother-bud axis. Maximum projection inverted images of cells expressing 3xGFP-Tub2 (DLY24515). Right: Each dot represents the angle of a spindle 5 s prior to anaphase onset in a single-budded cell (n=32 spindles). (B) Spindle orientation in multinucleate cells. Left: Maximum projection inverted images of cells expressing 3xGFP-Tub2. For a given spindle, illustrated with green asterisk, we measured two angles, one to each of the two mother-bud axes, 5 s prior to anaphase onset. Green denotes the angle to the axis of the bud that the spindle pole enters, and purple denotes the angle to the other axis. Right: Each dot represents one spindle in a multi-budded cell; green and purple dots report angles to different mother-bud axes as illustrated in the example. Mean angle is indicated with a bar (n=42 spindles). (C) Left: examples of cells that do (top) or do not (bottom) orient spindles along the mother-bud axis prior to anaphase. Time is in seconds. The oriented spindle (top) has a short astral microtubule extending towards bud-neck (green arrow). Maximum projection inverted time lapse images of cells expressing 3xGFP-Tub2 (DLY24515). Right: Angle between the spindle and the mother-bud axis during the 1 min before and after anaphase onset (time 0). (D) Graphs as in C for 20 additional multinucleate cells. Scale bars, 5 μm.

### Timing of nuclear entry into buds is variable

As metaphase spindles did not reliably pre-orient towards buds and anaphase was very rapid, we wondered whether entry of spindle poles into the buds occurred during or after anaphase. The time from anaphase onset to when a spindle pole entered a bud varied significantly. In mononucleate cells, spindle poles took 65 ± 24 s (n = 77 spindle poles) to enter a bud, while in multinucleate cells the timing was more variable and often took longer (86.9 ± 48.5 s, n = 125 spindle poles) (Fig. 5A). Entry times were variable even for spindle poles in the same cell (Fig. 5B). Many nuclei did not enter a bud until after anaphase: while 58% of spindle poles (n=86) that entered a bud did so during anaphase (spindle length < 6 μm), the other 42% entered a bud later, often apparently after disassembly of the anaphase spindle (Fig. 5C,D).

**Figure 5:**
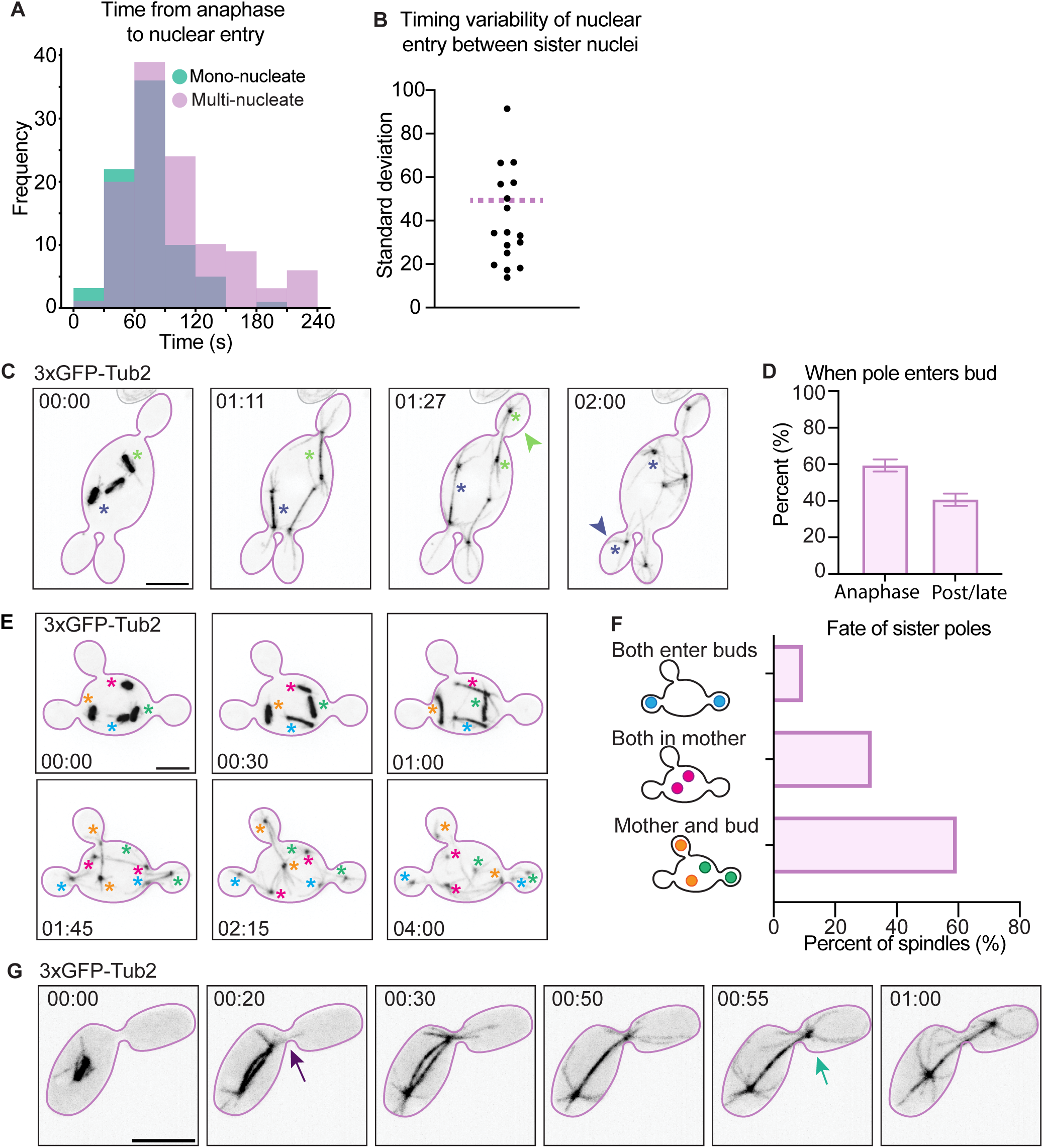
Nuclear entry into buds. (A) Interval between anaphase onset and nuclear delivery through the mother-bud neck, for uninucleate (green, n = 77 nuclei) and multinucleate cells (pink, n = 125 nuclei). Frequency indicates number of spindle poles. (B) Timing variability of nuclear delivery to sister buds. Each dot reports the standard deviation in s of nuclear entry times between sister-nuclei within the same cell (n=17 cells with 3 or more nuclei). The population standard deviation of nuclear entry for all buds from multinucleate cells is indicated by the purple dashed line. (C) Maximum projection inverted time lapse images of cell expressing 3xGFP-Tub2 (DLY24515) where spindle pole entry into bud occurs during (green arrowhead) and after (purple arrowhead) anaphase in the same cell. Asterisks indicate spindles (left panel) and spindle poles with colors matched to arrowheads. (D) Percent of wildtype spindle poles that enter buds during anaphase (when spindle poles are < 6 μm apart) or later (when spindle poles are > 6 μm apart and spindles may no longer be intact) (n = 86 spindle poles). (E) Maximum projection inverted time lapse images of multinucleate cell expressing 3xGFP-Tub2 (DLY24515). The 4 spindles and corresponding spindle poles are denoted with colored asterisks (blue, orange, green, red). (F) Quantification of sister pole destinations (n = 76 cells with 3 or more nuclei). Colored nuclei correspond to the spindle outcomes illustrated by the corresponding colors in (E). (G) Maximum projection inverted time lapse image of a cell expressing 3xGFP-Tub2 (DLY24515). An astral microtubule (purple arrow) extends through the bud neck into the bud before the attached spindle pole enters into a bud (green arrow). Astral microtubules penetrated the neck before the poles entered the buds in all cells observed (n = 30). Scale bars, 5 μm. Time in minutes: seconds.

Even within a single cell, different nuclei could enter buds during or after anaphase (Fig. 5C). These findings suggest that nuclear segregation can be effectively completed after spindle disassembly.

### The poles of a single spindle can vary in their fate

Given the lack of pre-orientation of spindles, we wondered if there was any pattern to the destinations of sister nuclei derived from a given spindle. In mononucleate single-budding yeasts, each spindle must deposit one nucleus in the bud and one in the mother cell, and that was also the case for mononucleate *A. pullulans* cells. However, in multinucleated cells we observed more variable outcomes. We tracked the destinations of sister poles from each spindle in cells with three or more nuclei. For 59% of spindles (n=76), one sister pole was delivered to a bud, and the other pole remained in the mother. However, for 32% of spindles both sister poles remained in the mother, and for 9% of spindles both sister poles entered buds (Fig. 5E,F). Therefore, it does not appear that sister nuclei have predetermined “fates”. Nevertheless, some mechanism must act to ensure that each bud receives at least one nucleus.

### Astral microtubules precede passage of spindle poles through the bud neck

The preceding characterization indicates that although mitotic spindles do not pre-orient towards buds before anaphase in *A. pullulans*, spindle poles are nevertheless delivered to every bud by about 5 min after anaphase onset. This suggests that there is a very rapid and effective mechanism for spindle poles to locate and enter each bud. As studies in *S. cerevisiae* indicated that astral microtubules guide the spindle pole to the bud, we asked whether nuclear entry into each bud in *A. pullulans* was preceded by astral microtubule contact with the bud. In the subset of metaphase spindles that did become oriented towards a bud, we often observed a short astral microtubule extending from one pole to a bud neck followed by the spindle moving closer to the neck (Fig. 4C). During and after anaphase, entry of a spindle pole into a bud was also preceded by penetration of an astral microtubule through that bud neck in every case we observed (n = 30 spindles) (Fig. 5G). These observations suggest that astral microtubules guide the spindle poles into the buds.

### Localization of dynein is consistent with a role in spindle pole positioning

How might astral microtubules draw the spindle poles into buds? In *S. cerevisiae,* astral microtubules are guided into the bud by the polarized actin cytoskeleton (Theesfeld et al., 1999; Hwang et al., 2003), and then spindle poles are pulled towards the bud by dynein (Yeh et al., 1995; Carminati & Stearns, 1997). To ask whether dynein might associate with astral microtubules at the bud cortex in *A. pullulans*, we tagged endogenous Dhc1, the dynein heavy chain, with three tandem copies of codon-optimized mCherry. Dynein tagging did not have any effect on colony growth (Fig. S3). In interphase budded cells, dynein puncta were present at bud tips and along microtubules (Fig. 6A). In metaphase cells, which lack long astral microtubules, dynein puncta were present around the cortex of both mother and buds (Fig. 6B). During anaphase, dynein was sometimes present along astral microtubules extending into the buds (Fig. 6C). In some cases, the tips of curved microtubules appeared to co-localize with dynein puncta in the bud (Fig. 6C). Later after spindle disassembly, dynein puncta were observed most brightly on both sides of the bud neck (Fig 6D). Thus, dynein puncta are present at astral microtubule contacts with the bud cortex, suggesting that dynein could potentially provide force to pull spindle poles into the buds.

**Figure 6:**
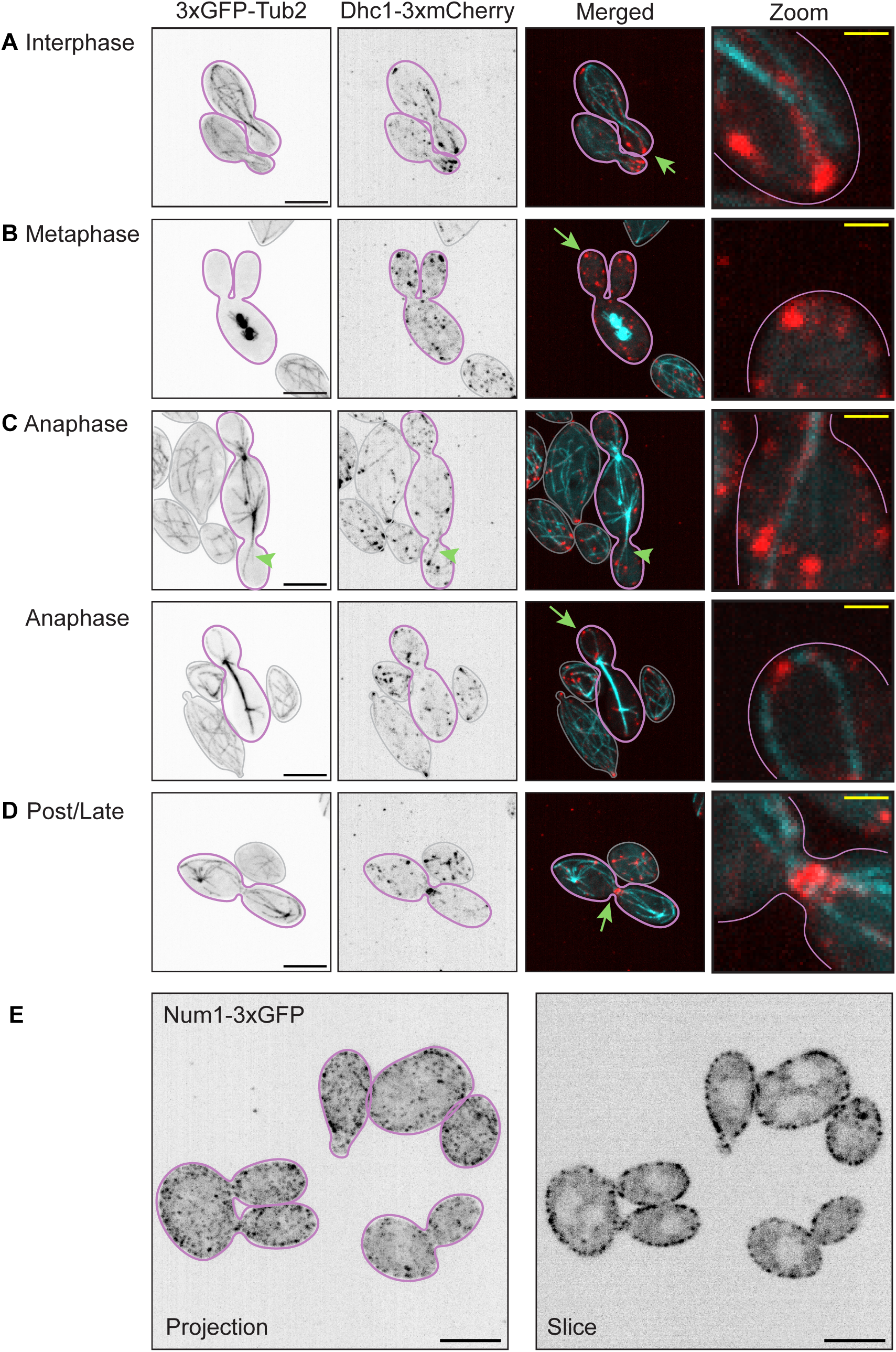
Dynein and Num1 localization. Inverted and merged maximum projection images of cells expressing 3xGFP-Tub2 and Dhc1-3xmCherry (DLY25434) illustrating distribution of dynein and microtubules in (A) interphase, (B) prophase and metaphase, (C) anaphase, and (D) late/post-anaphase cells. Green arrows (merged image) indicate magnified regions (right). (E) Maximum projection (left) and single slice (right) inverted images of cells expressing Num1-3xGFP (DLY25441). Num1 puncta are distributed around the cortex of both mothers and buds. Black scale bars, 5 μm; yellow scale bars, 1 μm.

To ask whether dynein contributes to nuclear segregation in *A. pullulans,* we attempted to generate *DHC1* null mutants. However, multiple attempts to knockout Dhc1 were unsuccessful, suggesting that dynein is essential for viability in *A. pullulans*.

### The dynein anchor Num1 is required for movement of nuclei into buds

Given that dynein was essential, we next looked at the localization and function of Num1, the dynein cortical anchor protein (Lee et al., 2003; Farkasovsky and Kuntzel, 2001; Markus and Lee, 2011). We tagged the endogenous *A. pullulans* Num1 with three tandem copies of codon-optimized GFP. Num1 tagging did not have any effect on colony growth (Fig. S3). Num1-3xGFP was concentrated in small puncta all along the cell cortex in both mothers and buds (Fig. 6E), consistent with Num1 functioning as a cortical anchor for dynein. We were able to generate Num1 viable deletion mutants, although they were very sick and grew very slowly (Fig. S3).

Remarkably, the majority of cells in *num1* mutants were anucleate and exhibited few microtubules (Fig 7A). Imaging of cells that did contain nuclei showed that they did make, and then elongate, mitotic spindles (Fig. 7B). However, in most cases, the divided nuclei stayed in the mother, with only 24% of buds receiving a nucleus (n=161 buds from 64 mothers)(Fig. 7B,C). Astral microtubules still entered buds and extended to bud tips, indicating that Num1 is not necessary for that step, but most spindle poles did not follow the attached astral microtubules into the buds (Fig. 7B,C, Supplemental Video 5). In the few cases where nuclei did enter buds, they almost always did so early during anaphase while the spindle was still intact (Supplemental Video 5). In some of these cases, the spindle pole exhibited a prolonged pause at the neck, unlike in wildtype cells (Fig. 7D, Supplemental Video 5). These observations suggest that the bud neck poses a mechanical barrier to passage of the nucleus and that although spindle-derived forces can push the nuclei through the neck, they often fail to do so.

**Figure 7:**
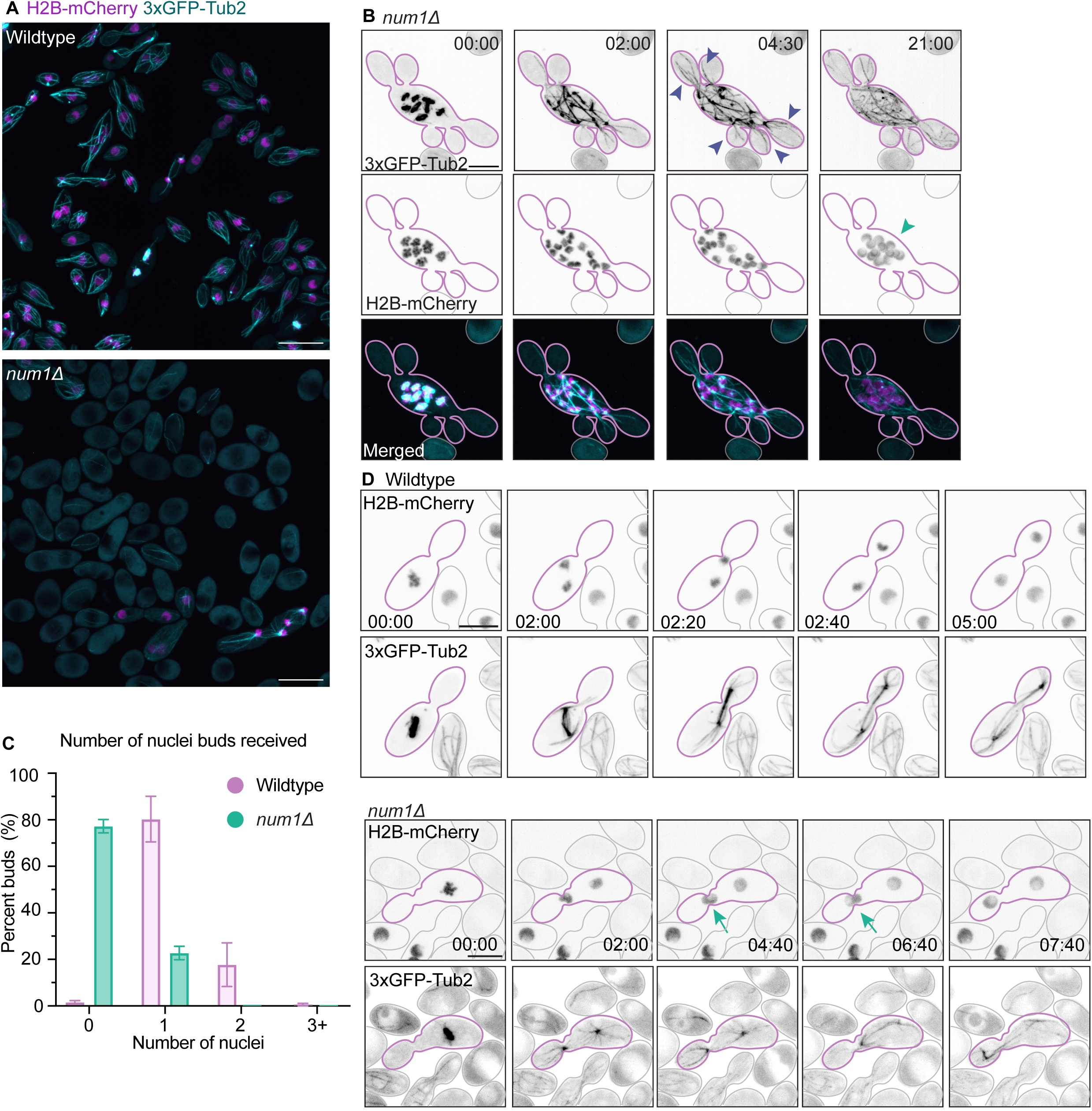
Nuclear inheritance in num1Δ cells. (A) Merged maximum projection images of wildtype (upper, DLY24568) and *num1*Δ (lower, DLY25357) cells expressing H2b-mCherry (magenta) and 3xGFP- Tub2 (blue). Many *num1*Δ cells lack nuclei and microtubules. (B) Maximum projection inverted and merged time lapse images of multinucleated *num1*Δ cell undergoing mitosis. Image intensity was scaled differently in inverted microtubule panels so that all microtubule features could be appreciated. For comparison of equally-scaled images see merged panel or Supplemental Movie 5. While spindles still produce astral microtubules that enter and extend to bud tips (purple arrowheads), the nuclei remain in the mother (green arrowhead) leaving buds anucleate after mitosis. (C) Number of nuclei inherited by each bud following mitosis in wildtype (pink, n = 1233 buds) and *num1*Δ (purple, n = 163 buds) cells. (D) Maximum projection inverted time lapse images of wildtype (upper) and *num1*Δ (lower) cells where buds do inherit a nucleus. Strains as in (A). Nuclei pass through the neck quickly in wildtype cells but pause at the neck in the *num1*Δ cell (green arrows). Black scale bars, 5 μm; white scale bars, 10 μm. Time in minutes: seconds.

For most nuclei, it appears that Num1, and by extension dynein, is needed to pull the nuclei through the neck into the buds.

### Nuclear movements in post-anaphase cells

Given the apparent microtubule-based pulling of nuclei into buds by cortical dynein, why don’t all nuclei go into buds? If pulling forces from dynein at the bud cortex drag spindle poles into buds, why are there any nuclei retained in the mothers? And why does each bud tend to get only a single nucleus? The observed segregation pattern suggests that in addition to bud-directed pulling forces, there must be other forces that prevent additional nuclei from entering already-occupied buds. Tracking nuclear movements with rapid time-lapse imaging (2.5 s intervals) showed that the centroids of the nuclear H2B-mCherry signals moved rapidly (0.1-0.6 μm/s) and erratically for >15 min following anaphase onset (Fig. 8A,B). Such movements could in principle be driven by pushing forces arising from contacts between growing astral microtubules and the cell surface or other objects. Focusing on post-anaphase cells (>2 min after anaphase onset), we found that nuclear movements slowed after the first few minutes (Fig. 8B), concomitant with the time when nuclei entered buds (Fig. 5A). Tracking nuclei in mothers and buds separately indicated that nuclei moved less once they were in buds (Fig. 8C), perhaps because the smaller bud size does not allow for longer excursions. If these rapid nuclear movements do arise from pushing forces, it could be that repulsion of later-arriving nuclei by the early-arriving nuclei explains why buds tend to acquire only one nucleus. It would also explain why larger buds are more likely to inherit two (see Figure 1G) as the first nucleus may move far enough away from the bud neck to permit the second to enter.

**Figure 8:**
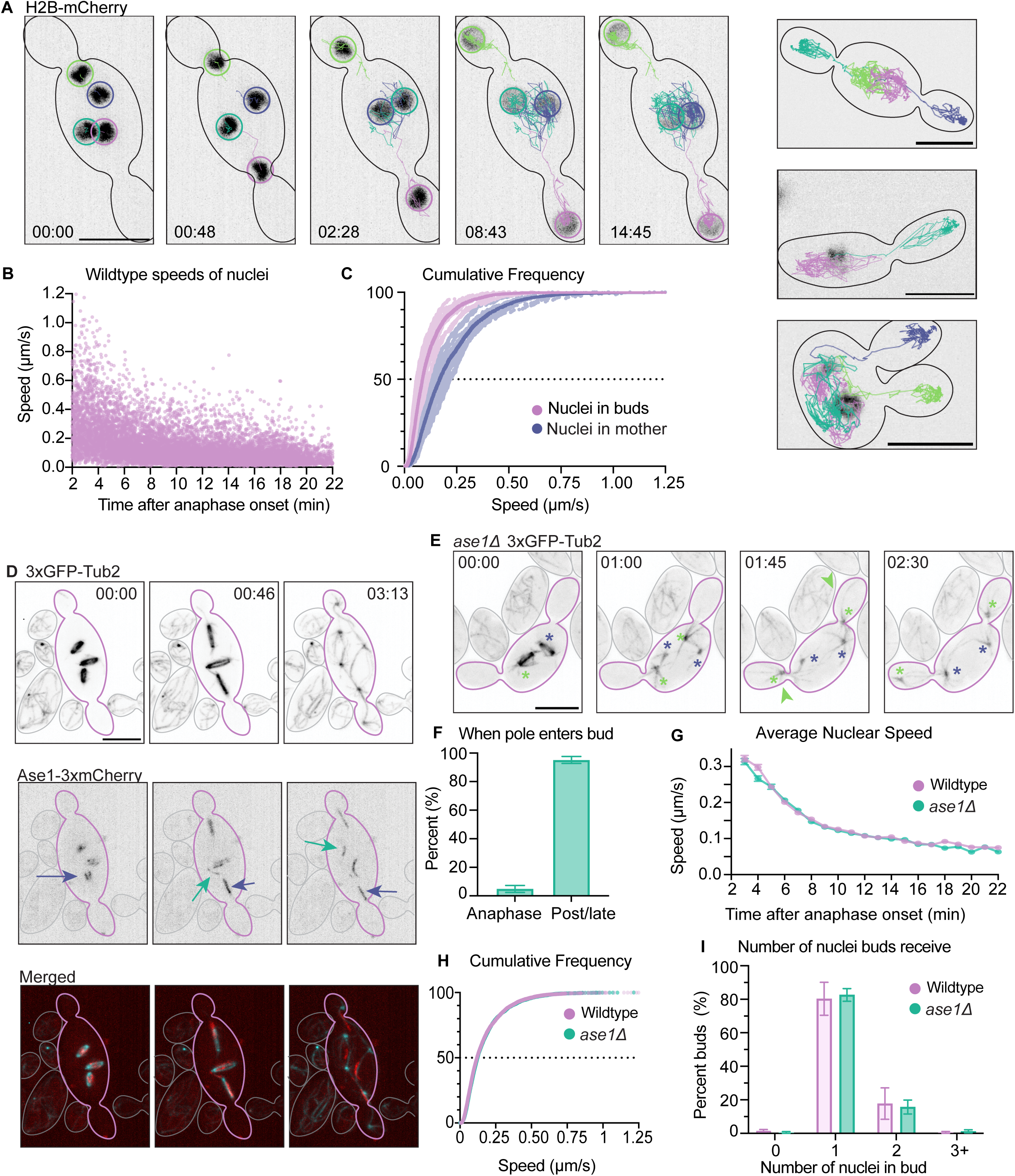
Ase1 role in nuclear segregation. (A) Left: Maximum projection inverted time lapse images of cells expressing H2B-mCherry (DLY24853) with overlay of centroid tracking where each post-mitotic nucleus’s movement in a different color. Right: nuclear tracks for three additional cells. (B) Instantaneous speeds from 26 nuclei in 10 wildtype cells (n = 12,468 speed measurements). (C) Cumulative frequency plot of nuclear speeds measured at 2-15 min after anaphase onset. Nuclei that are delivered to buds (pink, n = 14 nuclei) move less than those that remain in the mother (purple, n = 14 nuclei). (D) Maximum projection inverted and merged time lapse image of cells expressing 3xGFP-Tub2 and Ase1-3xmCherry (DLY25358). Ase1 accumulates at spindle midzones (purple arrows) and at overlap zones between astral microtubules (green arrows) during metaphase and anaphase. (E) Maximum projection inverted time lapse image of mitosis in an *ase1Δ* mutant cell expressing 3xGFP-Tub2 (DLY25382). Asterisks indicate spindles and spindle poles. Anaphase occurred entirely within the mother cell followed by the delivery of spindle poles from the green spindle into buds (green arrowheads). (F) Percent of *ase1Δ* spindle poles that entered buds during (while spindle is intact) or after (after spindle disassembly) anaphase (n = 109 spindle poles). (G) Average speed of post-anaphase nuclei (+/- SEM) in wildtype (pink, DLY24853) and *ase1Δ* (green, DLY25382) cells. (H) Cumulative frequency plot of nuclear speeds (as in C) for wildtype (pink, n = 8,100 speeds from 26 nuclei) and *ase1Δ* (green, n = 6814 speeds from 24 nuclei) cells. (I) Number of nuclei inherited by each bud following mitosis in wildtype (pink, n = 1233 buds) and *ase1Δ* (green, n = 1195 buds) cells. Scale bars, 5 μm. Time in minutes: seconds.

A precedent for inter-nuclear repulsion comes from studies in multinucleate animal cells, such as the *Drosophila* syncytial blastoderm, where such forces are responsible for the even distribution of nuclei within the syncytium. In that system, repulsive forces between asters arise from antiparallel overlap of astral microtubules (Deshpande et al., 2021). Overlap zones are stabilized by the crosslinker Prc1/Ase1, which allows kinesin IV and kinesin V motors to slide the microtubules apart. A similar situation applies to the central spindle of mitotic yeast cells, where Prc1/Ase1 cross-links antiparallel microtubules from each spindle pole, allowing kinesin V to drive the poles apart (Yamashita et al., 2005; Juang et al., 1997). These findings prompted us to ask whether Ase1 plays a similar role in nuclear distribution in *A. pullulans*.

### Localization of the microtubule crosslinker Ase1 in *A.pullulans*

To visualize where Ase1-stabilized antiparallel microtubules might occur in *A. pullulans*, we tagged the endogenous Ase1 with 3 tandem copies of codon-optimized mCherry. Tagging did not have an effect on colony growth (Fig. S3). Ase1 was localized to spindle midzones during metaphase and early anaphase (Fig. 8D). However, during and after anaphase, Ase1 also decorated presumed antiparallel microtubule bundles between poles that originated from different spindles (Fig. 8D). This suggests that astral microtubules from different poles interact with each other during and after anaphase, which could lead to repulsive forces between asters that prevent additional nuclei from entering buds once the first nucleus has been drawn in.

### Mitosis and nuclear segregation in cells lacking Ase1

To assess potential roles for antiparallel microtubule bundles in nuclear segregation, we generated *A. pullulans ase1* deletion mutants. *ase1* mutants were viable and, unlike *num1* mutants, had no detectable colony growth defect compared to wildtype (Fig. S3). In *ase1* mutants, anaphase spindles broke down at a shorter length (3.5 ± 0.9 μm, n = 118 spindles), and we did not observe any spindles reach 6 μm. This finding is consistent with a role for Ase1 in stabilizing interpolar microtubule bundles during anaphase. At the time of spindle breakdown, both spindle poles were still in the mother in 95% of cells (n = 247 spindle poles)(Fig. 8E,F).

Nevertheless, spindle poles remained highly dynamic and entered buds after anaphase (Fig. 8E, Supplemental Video 6). Tracking nuclear movements in *ase1* mutants, we detected no difference in the distribution of speeds compared to wild-type cells (Fig. 8G,H). Thus, Ase1- mediated antiparallel microtubule interactions between asters are not necessary for post-anaphase nuclear movements.

Despite the premature spindle breakdown, we did not observe any change in the frequency with which buds inherited different numbers of nuclei in the *ase1* mutants (Fig. 8I). Thus, post-anaphase nuclei have an effective mechanism to find and enter buds, and they can also prevent multiple nuclei from entering a single bud, even in the absence of Ase1.

## Discussion

### Mitosis in *A. pullulans*

Here we describe the dynamics of mitosis in the multinucleate and multibudding yeast *A. pullulans*. Fungi exhibit considerable variety in their patterns of mitosis: mitosis may be rapid or slow, closed or semi-open (De Souza et al., 2004; Theisen et al., 2008; De Souza and Osmani, 2009; Aoki et al., 2011), synchronous or asynchronous (Clutterbuck, 1970; Serna and Stadler, 1978; Gladfelter et al., 2006; Gladfelter, 2006). *A. pullulans* exhibits a rapid, synchronous, and semi-open mitosis that is most similar to that described for the filamentous fungus *A. nidulans*, and quite different from that described for most budding yeasts. Our findings on live cells with a fluorescent tubulin probe are fully consistent with an earlier study using immunofluorescence to examine tubulin distribution in the hyphal form of *A. pullulans* (Kopecka et al., 2003).

Interphase cells contain an extensive network of cytoplasmic microtubules that fills the mother cell and extends microtubule plus ends into each growing bud. At the onset of mitosis, the interphase network is dismantled within about 1.5 min, concomitant with formation of a short bar-shaped spindle in each nucleus. During the brief 4 min metaphase, almost all of the tubulin signal is in the nuclear spindles and there are only a few very short astral microtubules. This changes rapidly and dramatically during anaphase. Spindle elongation occurs over about 1.5 min and coincides with a loss of tubulin from the spindle, accompanied by the rapid growth of a robust cytoplasmic microtubule aster extending from each spindle pole. Nuclei and asters remain highly mobile during an extended post-anaphase period of about 15 min before cytokinesis. During and after anaphase, nuclei are delivered to every bud, with errors leading to anucleate buds being very rare. Our experiments begin to explain how *A. pullulans* cells can achieve such accurate nuclear segregation in the face of the geometric and mechanical challenges posed by the multi-budded cell morphology.

### Potential mechanisms of nuclear segregation in *A. pullulans*

We initially considered two potential nuclear segregation mechanisms, inspired by findings in budding yeasts and hyphal fungi. First, as in the yeasts *S. cerevisiae* (Segal and Bloom, 2001), *Cryptococcus neoformans* (Chatterjee et al., 2021), *Candida albicans* (Martin et al., 2004), and *Ustilago maydis* (Fink et al., 2006), segregation might be driven by the mitotic spindles. In those phylogenetically diverse but uninucleate yeasts, the spindle is pre-oriented along the mother-bud axis and spindle elongation provides much of the force required to squeeze a nucleus through the narrow mother-bud neck. Second, as in the multinucleate hyphae of *Ashbya gossypii* (Anderson et al., 2013), *Neurospora crassa* (Plamann et al., 1994), and *A. nidulans* (Xiang et al., 1994), nuclei might become evenly distributed due to cytoplasmic microtubule-driven interactions between neighboring nuclei.

The multibudded geometry of *A. pullulans* presents formidable challenges for each of these potential nuclear segregation mechanisms. For spindle-driven segregation, it is not obvious how the different spindles in multinucleated cells would coordinate their actions to distribute their nuclei evenly to every bud. For nuclear repulsion-driven segregation, it is not obvious how the nuclei would find and then squeeze through the narrow neck apertures to deliver the nuclei into buds. Our findings are incompatible with both starting hypotheses, but they suggest that a combination of mechanisms derived from the yeast and hyphal systems might suffice to yield accurate segregation in this challenging geometry.

### Spindle forces are not required to deliver nuclei into buds

As noted above, metaphase spindles in *A. pullulans* lacked extensive astral microtubules. Therefore, it was perhaps unsurprising that prior to anaphase, spindles did not orient along the mother-bud axis. Even during anaphase, spindles often failed to align or deliver nuclei into buds prior to spindle breakdown. Most strikingly, accurate nuclear segregation was preserved in *ase1* mutants, in which anaphase spindles almost always (95%) broke down prior to entry of a daughter nucleus into the bud. Thus, although spindles may assist in nuclear segregation, post-anaphase *A. pullulans* cells have an effective nuclear segregation pathway that can deliver nuclei into buds without assistance from the mitotic spindles.

### Post-anaphase nuclear motility

Inter-aster repulsive interactions are thought to result in even spreading of nuclei in multinucleate animal cells (e.g. *Drosophila* syncytial blastoderm) and could underlie the even spreading of nuclei in filamentous fungi (Anderson et al., 2013; Deshpande et al., 2021). One documented mechanism for such repulsive interactions involves kinesin-driven sliding apart of antiparallel microtubules bundled by the cross-linker Prc1/Ase1 (Deshpande et al., 2021). In *A. pullulans,* we found that individual nuclei associated with microtubule asters remained highly mobile for several minutes after anaphase and that Ase1-decorated overlap zones could form between asters. However, Ase1 was not required for post-anaphase nuclear movements or for normal nuclear segregation into buds. The mechanism that drives post-anaphase movement remains to be determined but may include pushing of astral microtubules against the cell cortex and/or dynein-mediated sliding of astral microtubules along the cortex.

### Num1-mediated pulling forces deliver nuclei into buds

If inter-nuclear repulsive forces were responsible for delivering post-anaphase nuclei into buds, then astral microtubules would not need to enter buds prior to nuclear segregation.

However, we found that entry of a nucleus into a bud in *A. pullulans* was invariably preceded by entry of one or more associated astral microtubules into the bud. This suggests that as in mononucleate yeasts (Miller and Rose, 1998; Hwang et al., 2003), there is a pathway to guide astral microtubules into buds. The nature of that pathway remains to be determined.

As dynein-mediated pulling has been shown to contribute to nuclear entry into buds of mononucleate yeasts (Yeh et al., 1995), we attempted to assess the role of dynein in nuclear segregation in *A. pullulans*. In *S. cerevisiae*, dynein is delivered to the tips of astral microtubules in an inactive form and then off-loaded and activated at cortical sites enriched for the dynein anchor Num1 (Markus and Lee, 2011). Num1 is a fungal analogue of the animal protein NuMA (Greenberg et al., 2018), and we found that as in other yeasts, Num1 was localized to patches throughout the cortex of mothers and buds in *A. pullulans*. Dynein was localized to both astral microtubules and cortical patches, consistent with the idea that dynein anchored to cortical Num1 might drive sliding of astral microtubules along the cortex and thereby contribute to nuclear motility.

We were unable to recover viable dynein deletion mutants, but we did recover mutants lacking Num1. *num1* mutants exhibited severe growth defects associated with the production of large numbers of anucleate cells. When *num1* mutant cells that did contain nuclei went through mitosis, they formed intranuclear spindles and elaborated astral microtubules in anaphase that efficiently extended to the tips of buds. Anaphase proceeded with similar timing as in wild-type cells and nuclei exhibited rapid post-anaphase motility. However, the nuclei in *num1* mutants only rarely penetrated through the bud necks. These findings suggest that Num1/dynein-derived forces are needed to efficiently pull the nuclei through the narrow mother-bud necks.

The minority of nuclei that did enter buds in *num1* cells usually did so during anaphase, suggesting that spindle-derived forces are also capable of pushing the nuclei through the neck. We speculate that they often fail to do so because the spindles are not oriented towards the bud early enough in anaphase. Moreover, even when nuclear delivery to the bud succeeded, there appeared to be a delay with the nucleus stuck in transit at the neck, consistent with the hypothesis that most of the force that enables rapid transits of nuclei into buds in wild-type cells is provided by Num1/dynein.

The large number of anucleate cells observed in *num1* mutants suggests that anucleate buds can undergo cytokinesis and detach from the mother cell. This may indicate that *A. pullulans* cells do not have a checkpoint analogous to the spindle position checkpoint described in *S. cerevisiae* (Yeh et al., 1995; Bloecher et al., 2000; Pereira et al., 2000; Wang et al., 2000).

### Mechanisms of nuclear segregation in a multibudding yeast

Based on our findings, we suggest a pathway that can yield accurate nuclear inheritance in multi-budded *A. pullulans*. During metaphase, the large majority of tubulin is recruited to the intranuclear spindles, leaving very little in the cytoplasm. Lacking cytoplasmic microtubules, the spindles are untethered from the cortex and do not orient towards buds. During anaphase, many spindle microtubules disassemble, and the tubulin released from the nucleus drives rapid growth of cytoplasmic microtubule asters. Astral microtubules are efficiently directed into buds, where they interact with dynein and Num1 to pull the attached spindle poles into the buds. Entry of one nucleus into a bud then discourages subsequent entry of other nuclei. We speculate that the presence of one cytoplasmic aster in a bud may bind and obscure the limited available Num1/dynein cortical sites in the bud, so that asters from nuclei still in the mother are unlikely to be pulled into an occupied bud. This would explain why most buds receive just one nucleus.

This description is plausible but leaves several open questions. How are astral microtubules directed into the buds? Are there nuclear repulsion pathways independent of Ase1? Do Num1 patches in the mother cell also contribute to nuclear motility? If so, how do the bud-localized patches assure that nuclei override mother-localized pulling forces to drive nuclei into buds? Further studies into mitosis mechanisms in the challenging multibudded geometry will be illuminating with regard to these and other questions.

## Methods

### *Aureobasidium pullulans* strains

All *Aureobasidium pullulans* strains are in the *EXF-150* strain background (Gostincar et al., 2014). Genetically-encoded probes and mutations were introduced into *A. pullulans* by means of chemical transformation (Wirshing et al., 2024) using homologous recombination strategies with vectors or PCR products as described (Colarusso et al., 2025). Strains are listed in Table S1. For each strain, 3 separate transformants were checked for consistent phenotypes, growth rates, and protein localization. Either hygromycin or nourseothricin resistance genes were used for selection of gene deletions. For fluorescently tagged proteins, if an additional copy was expressed, it was from the *Saccharomyces cerevisiae ACT1* promoter except for histones which were expressed from the chytrid H2AB promoter as described (Petrucco et al., 2024). Fluorescent probes were integrated at the *URA3* locus as described (Wirshing et al., 2024). GFP and mCherry tags were designed so that each amino acid was encoded with its most frequently used codon. Three GFPs or mCherrys were used in tandem to obtain brighter signals (Colarusso et al., 2025). To determine the effects of genetic alterations on growth and colony morphology, serial dilutions containing 5000, 500, 50, and 5 cells were spotted onto the same YPD plate and incubated for 2 d at 25°C.

Proteins were identified by BLASTp (protein-protein BLAST) using *Saccharomyces cerevisiae (S.c.)* proteins as query sequences. All *A. pullulans* proteins in this study were the best hit. The *A. pullulans* proteins were then checked on JGI Mycocosm to ensure they contained the same PFAM domains as the *S.c*. homologs. A list of proteins used in the study are in Table S2.

### Cell growth and imaging conditions

Yeast cells were grown in at room temperature (21-23°C) in YPD medium (2% dextrose [VWR], 2% bacto-peptone [Gibco], 1% yeast extract [Gibco], and for plates 2% difco-agar [Becton Dickinson and Company].

To culture yeast for imaging experiments, cells were grown overnight at 25°C in YPD to a density of 6-8 x 10^6^ cells/mL. 400 μL of cells were pelleted at 9000 x g for 10 s and resuspended in 100 μL of YPD. 2-4 μL of cell suspension was placed in an ibidi 8 well glass bottom μ-slide with a 200 μL slab of complete synthetic media, (CSM: 2% dextrose [VWR], 0.7% yeast nitrogen base without amino acids [VWR], and 0.08% complete supplement mixture [MP Biomedicals]) on top, solidified with 2% agarose [Fisher]. All imaging experiments were conducted at room temperature (21-23°C).

### Live cell imaging

To measure brightness of GFP tagged Tub2 with 1x, 2x, or 3x GFP, cells were imaged on a widefield Nikon ECLIPSE Ti2 inverted microscope with a Plan Apo 60x Oil Immersion Objective (NA 1.40; Nikon Instruments), a sCMOS pco.edge camera (Excelitas Technologies), and a X-Cite XYLIS LED Illumination System (Excelitas Technologies) controlled by NIS-Elements software (Nikon Instruments). The entire cell volume was acquired using 13 Z-slices at 0.5 μm step intervals. Exposure times of 200 ms at 5% laser power (excitation 488 nm) were used.

To monitor the dynamics of Bim1-3xmCherry on the tips of growing microtubules, cells expressing Tub2-3xGFP and Bim1-3xmCherry were imaged on a Zeiss 980 scanning confocal microscope using a 63x oil objective with N.A. 1.4 in confocal mode. Images of GFP and mCherry probes were acquired simultaneously by excitation with 488 nm and 561 nm lasers. 45 frames were acquired at a rate of one frame every 1.878 seconds.

To measure bud-neck widths, cells expressing Cdc3-tdTomato were imaged on a Zeiss Axio Observer widefield microscope using a 63x oil objective, and a Hamamatsu Flash 4 camera. The entire cell volume was acquired using Z-slices at 0.32 μm step intervals. Exposure times of 500 ms at 21% laser power (excitation 561 nm) were used.

Time-lapse images to count the number of nuclei in buds were acquired with an Andor Revolution XD spinning-disk confocal microscope (Andor Technology) with a CSU-X1 5000- rpm confocal scanner unit (Yokogawa) and a UPLSAPO 60x/1.3 silicon oil objective (Olympus) controlled by MetaMorph software (Molecular Devices). Images were captured by iXon Life 888 EMCCD camera (Andor Technology). The entire cell volume was acquired using 14 Z-slices at 0.8 µm steps. To image H2B-tdTomato, H2B-mCherry or NLS-tdTomato probes, we used exposure times of 200 ms at 8-15% laser power (excitation 561). DIC images were acquired with an exposure time of 50 ms.

Imaging of all other fluorescent probes, including images that include nuclei and another fluorescent probe, were acquired with a Nikon Ti2-E stand equipped with a Yokogawa CSU-W1 spinning disk confocal unit and a 100x/1.35 silicone oil objective. Images were captured using a Hamamatsu ORCA-Fusion BT sCMOS camera controlled by NIS-Elements software (Nikon instruments). To capture the entire volume of the cell, 8-24 Z-slices were acquired at 0.48 µm or 0.8 µm for only nuclei. To image cells expressing 3xGFP-Tub2 an exposure time of 200 ms at 5- 7% laser power (excitation 488) was used. To image cells expressing Pak1-GFP an exposure time of 500 ms at 5% laser power (excitation 488) was used. To image Bim1-3xmCherry, Ase1- 3xmCherry, or H2B-mCherry an exposure time of 200 ms at 10-25% laser power (excitation 561) was used. To image H2B-mCherry rapidly for 3D-tracking of nuclei, an exposure time of 100 ms at 10% laser power was used. For two-color imaging, the full red z-stack was taken followed by the full green z-stack. This led to a delay of up to 3.4 s between the red and green images which sometimes led to misalignment between the channels when objects were moving rapidly, as with histone and Tub2 signals during anaphase. Time lapse images were acquired at intervals between 5 s and 30 s as indicated. At least three independent movies were collected for each strain to visualize localization of fluorescent proteins.

### Image analysis

All image analysis was done in FIJI (Schindelin et al., 2012) unless indicated.

#### Scoring bud size and nuclear inheritance by sister buds

To unambiguously identify sister buds, we only scored those buds that formed simultaneously from the same mother cell during time lapse imaging. Number of nuclei inherited by each bud was scored visually. Bud volume was estimated from single plane DIC images assuming an ellipsoid shape with measured long and short axes.

#### Timing of post-anaphase interval

To estimate the timing of cytokinesis, time-lapse Z-stack images of cells expressing H2B-mCherry and Pak1-GFP were used. Mitosis onset was scored when histones condensed while cytokinesis was scored when Pak1 localized at the neck.

#### 3D Tracking of nuclei

To track nuclear movement after anaphase, 2.5 s interval time-lapse Z-stack images of wildtype and *ase1Δ* cells expressing H2B-mCherry were used. Nuclei were tracked using TrackMate (Ershov et al., 2022) through FIJI using the DOG detector, estimated object diameter of 1.8 µm, and simple LAP tracker. The centroid of the histone signal was tracked, although this was often skewed from the visual center of the nucleus due to the lower histone signal in the nucleolus. Manual editing of the tracks was sometimes necessary to ensure that the entire path of each nucleus was measured. The output of tracking was X,Y,Z, and time coordinates for the centroid of each nucleus. These coordinates were used to calculate instantaneous speeds.

Outlier speeds greater than 4 μm/s (n = 8 of 12468 speeds measured) were removed from the data set before analysis. Cumulative frequency plots were generated using data from between 2 and 15 min post anaphase onset, called visually based on when condensed chromosomes began to divide. For wildtype nuclei, the destination of each nucleus (mother or bud) was documented by eye.

#### Bud-neck width

To measure bud-neck width, Z-stack images of cells expressing Cdc3-tdTomato were used. The septin-ring width was measured from the medial z-plane where the septin width was widest, scoring cells with medium-sized buds.

#### Brightness of Tub2 probes

To quantify probe brightness, average intensity projections were generated. Background (signal outside of cells) was subtracted from each image and the signal intensity inside cells was measured.

#### Bim1 dynamics

To track Bim1-3xmCherry on the tips of growing microtubules, cells expressing Tub2- 3xGFP and Bim1-3xmCherry were imaged. To construct kymographs, we first fit curves to the image channels corresponding to microtubules using the Jfilament plugin for ImageJ, with the following parameters: stretch=100, alpha=20, beta=10, and gamma=400. Curves generally tracked microtubules well through time, although manual correction was at times necessary, e.g. when filaments intersected. Kymographs were produced from the intensity of pixels along these curves, using the Boundary Kymograph function provided by Jfilament, for both the Tub2 channel and Bim1 channel images. Microtubule kymographs were false-colored cyan while Bim1 kymographs were false-colored red, and the two kymographs were merged into a composite RGB image.

#### Microtubule behavior and dynamics

For all measurements of spindle and microtubule behavior, time-lapse Z-stack images of cells expressing 3xGFP-Tub2 were used. The distance between spindle poles was measured from the X, Y, Z position of each spindle pole. The number of astral microtubules a spindle had was determined by counting the number of astral microtubules from each individual pole and adding them together. We counted astral microtubules as those that emanated from either spindle pole independent from the bright, thick pole-to-pole spindle.

Astral microtubules were often curved, and their length was estimated by tracing the path on a maximum intensity projection 2D image. This would lead to some underestimation of the absolute astral length, due to extension along the Z dimension, but should not affect our conclusion that astral length increased as anaphase progressed.

For analyses from snapshot images, cells with spindle shorter than 3 µm were considered to be in metaphase, and those with spindle between 3-6 µm were considered to be in anaphase. Because the spindle intensity decreased as spindles lengthened, it was not obvious whether or not the spindles had disassembled when poles were further than 6 µm apart in 3D, and cells at that stage were classified as late/post anaphase. For analyses from movies, the time it took for spindle assembly was measured from the time a spindle first became visible to the time when the interphase microtubule network had disassembled. Metaphase was measured from the time no interphase microtubules present until the start of spindle elongation. Anaphase was measured from the start of spindle elongation until spindles reached 6 µm in length. 6 µm length was chosen because in most cases the longest structures we could confidently identify as an intact spindle were ∼6 µm in length.

To determine whether spindle elongation occurred synchronously, the length of each spindle in cells with two or more spindles was measured as described above every 30 s for 4.5 min or until the spindle was clearly disassembled.

Spindle orientation was measured as the angle between the spindle and the mother-bud axis. The mother-bud axis was determined as the straight line from the center of the bud tip through the middle of the bud neck. Measurements were taken on maximum intensity projection images 5 s prior to the onset of spindle elongation. For multi-nucleate cells the orientation of each spindle to every mother-bud axis was measured. To measure spindle orientation over time, the orientation of the spindle in mononucleate single-budded cells was measured every 5 s for 1 min before and after anaphase onset.

The fate of sister poles (i.e. whether they ended up in the mother, or in buds, or in mother and bud) was determined by tracking the destinations of the two spindles poles during mitosis.

The timing of nuclear entry was scored as the time from when spindle elongation started to when the spindle pole entered the bud. This was measured both mononucleate and multi- nucleate (2 or more nuclei) cells. The standard deviation of nuclear entry times between sister- nuclei was measured for cells with 3 or more buds. To determine whether a spindle pole entered a bud during or after anaphase, we scored whether spindles had elongated to < 6 µm (enter during anaphase) or > 6 µm (enter late/post-anaphase) at the time when the pole crossed the mother-bud neck. For *ase1Δ* cells, spindles disassembled before reaching 6 µm in length. If a spindle pole entered a bud when the spindle was intact it was called to enter during anaphase, and otherwise it was called to enter during late/post anaphase.

## Acknowledgements

We thank Clara Fikry and Yiqiao Zheng for comments on the manuscript. Some imaging was performed with instruments of the Duke Light Microscopy Core Facility, and we thank Lisa Cameron and Yasheng Gao for assistance with microscopy. This work was funded by NIH/NIGMS grant R35GM122488 to DJL, and by NSF grant MCB-2016022 to ASG.

---

**Video 1: Histones and Tubulin throughout mitosis.** Examples of cells with varying number of nuclei or buds throughout mitosis. During mitosis, each nucleus has one bar-shaped spindle. As the spindle elongates filaments (astral microtubules) grow and extend from each pole. Spindles become fainter and thinner as they elongate, and after spindle disassembly each nucleus is associated with one spindle pole that has microtubules emanating from it. Merged maximum projection time lapse images H2B-mCherry (magenta) and 3xGFP-Tub2 (cyan) (DLY 24568) at 20 s (left) or 25 s (right) intervals. Scale bar, 5 μm.

**Video 2**: **Bim1 puncta decorate the tips of astral microtubules**. Bim1 (EB1) puncta decorate the tips of astral microtubules. Merged maximum projection time lapse images of Bim1- 3xmCherry (red) and 3xGFP-Tub2 (cyan) (DLY 25139) at 4.4 s intervals. Scale bar, 5 μm.

**Video 3: Assembly of spindle and disassembly of interphase microtubule network**. The spindle forms while the interphase microtubule network disassembles. Inverted maximum projection time lapse images of 3xGFP-Tub2 (DLY 24515) at 5 s intervals. Scale bar, 5 μm.

**Video 4: Astral microtubules remain dynamic after anaphase**. As the spindle elongates, it gets lighter and thinner. After spindle disassembly the two spindle poles remain highly dynamic. Inverted maximum projection time lapse images of 3xGFP-Tub2 (DLY 24568) at 2.4 s intervals. Scale bar, 5 μm.

**Video 5: nuclear inheritance in *num1Δ*.** Most buds do not receive a nucleus, but when a nucleus does enter a bud, the spindle is usually intact. Maximum projection time lapse images of H2B-mCherry (magenta) and 3xGFP-Tub2 (cyan) (DLY 25357) at 25 s intervals. Scale bar, 5 μm.

**Video 6: *ase1Δ* cells undergo anaphase within the mother cell**. Anaphase takes place within the mother cell, then after spindle disassembly nuclei move into buds. Inverted maximum projection time lapse images of 3xGFP-Tub2 (DLY 25382) at 15 s intervals. Scale bar, 5 μm.

**Supplemental Figure 1:**
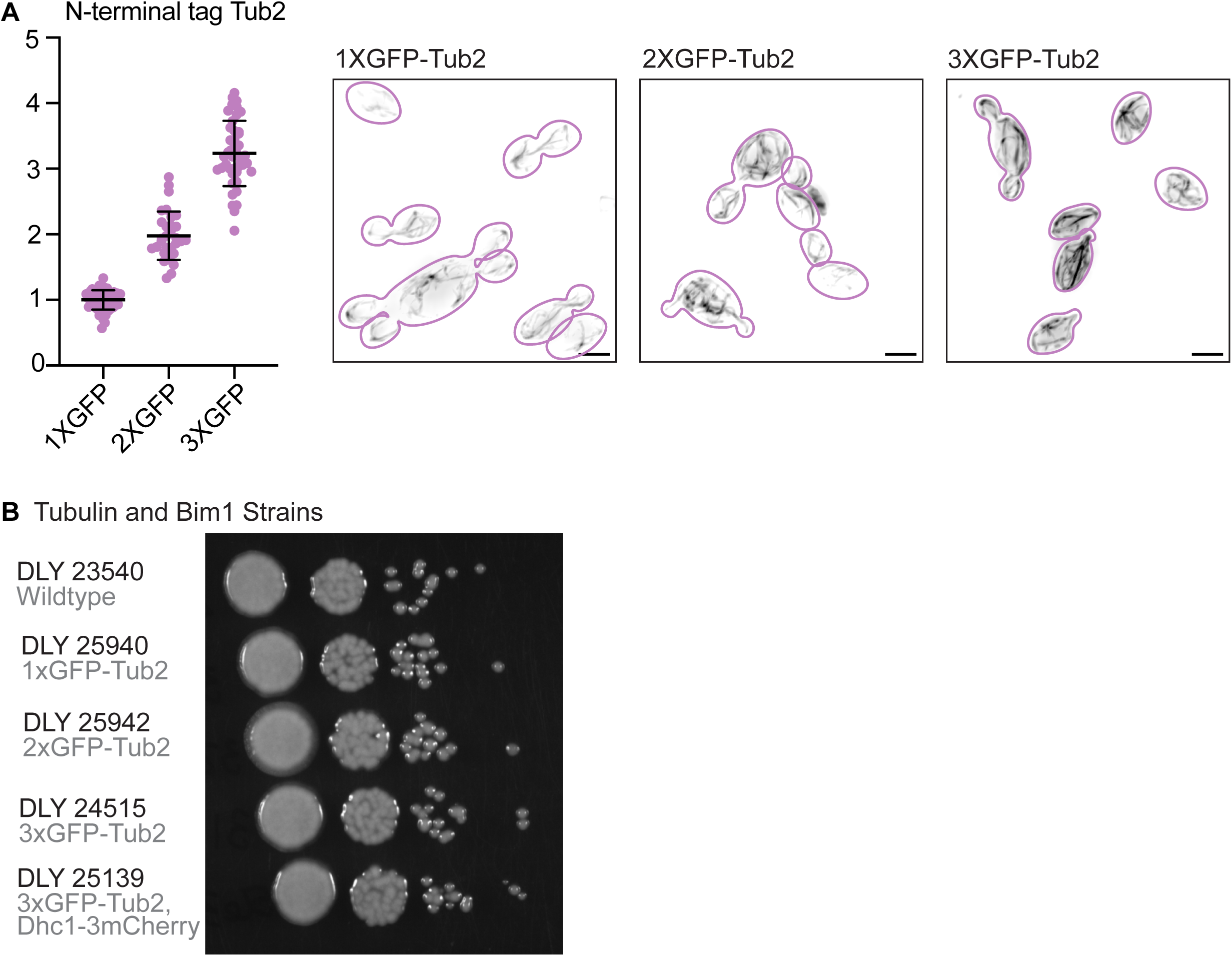
Visualizing microtubules in *A. pullulans*. (A) Tandem GFP probes enhance microtubule detection. Left: relative GFP signal per cell (n = 135 cells) for strains expressing GFP-Tub2 (DLY25940), 2xGFP-Tub2 (DLY25942), and 3xGFP-Tub2 (DLY24515). Right: Maximum projection inverted images of cells from the same strains, scaled equally. Scale bars, 5 μm. (B) Cells expressing 3xGFP-Tub2 and/or Bim1-3xmCherry do not exhibit colony growth defects. Cells of the indicated strains (5000, 500, 50, and 5 cells) were spotted onto YPD and grown for two days at 25°C.

**Supplemental Figure 2:**
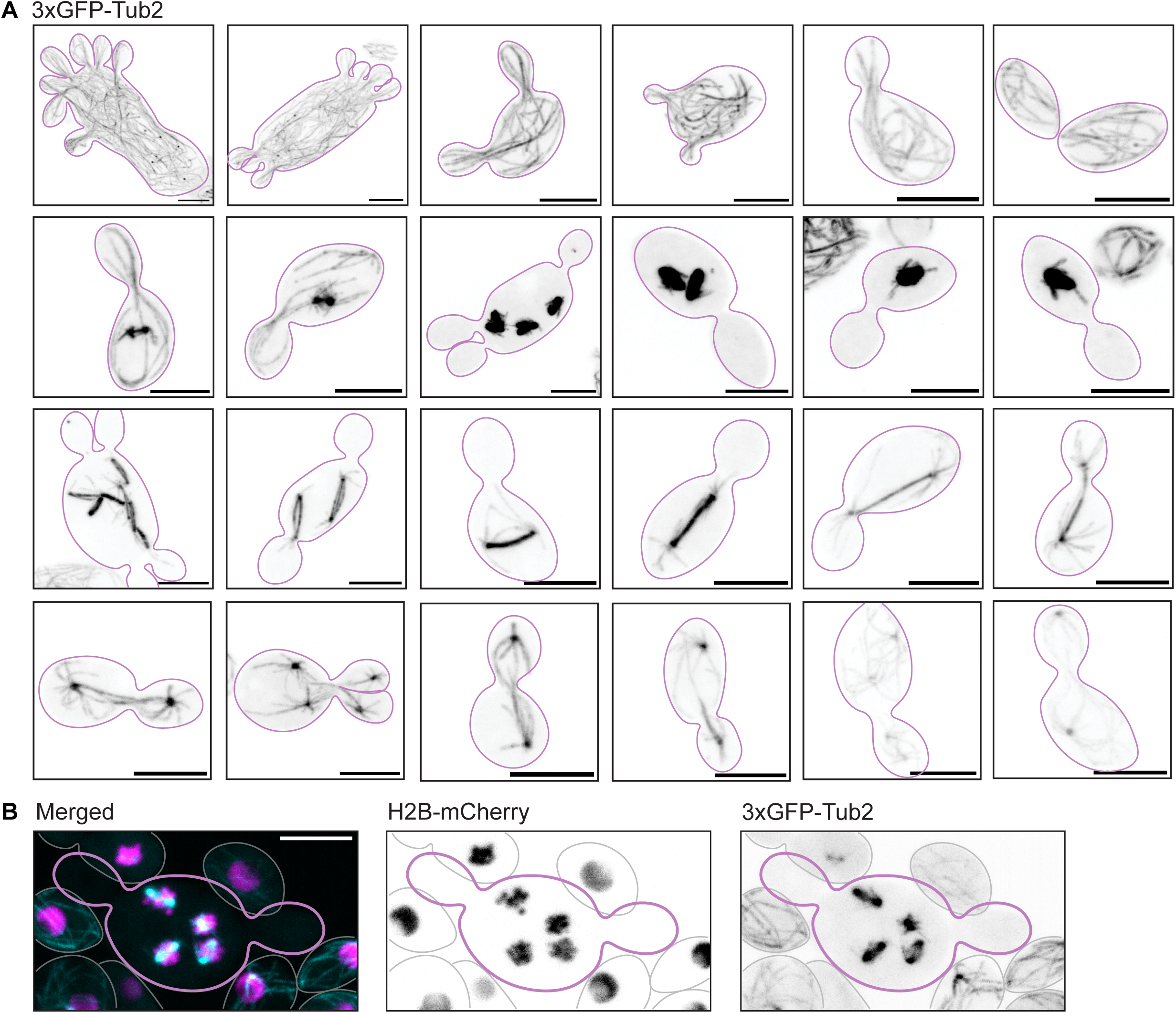
**Microtubules in *A. pullulans.*** (A) Inverted maximum projection images of cells expressing 3xGFP-Tub2 (DLY24515). These are the same cells as in Fig. 2A-D, but all images are scaled equally, showing that short spindles are much more densely packed with microtubules than anaphase spindles, which become fainter as they elongate. (B) One spindle is built for each nucleus. Maximum projection inverted and merged image of cell expressing 3xGFP-Tub2 and SpH2B-mCherry (DLY24568). Scale bars, 5 μm.

**Supplemental Figure 3:**
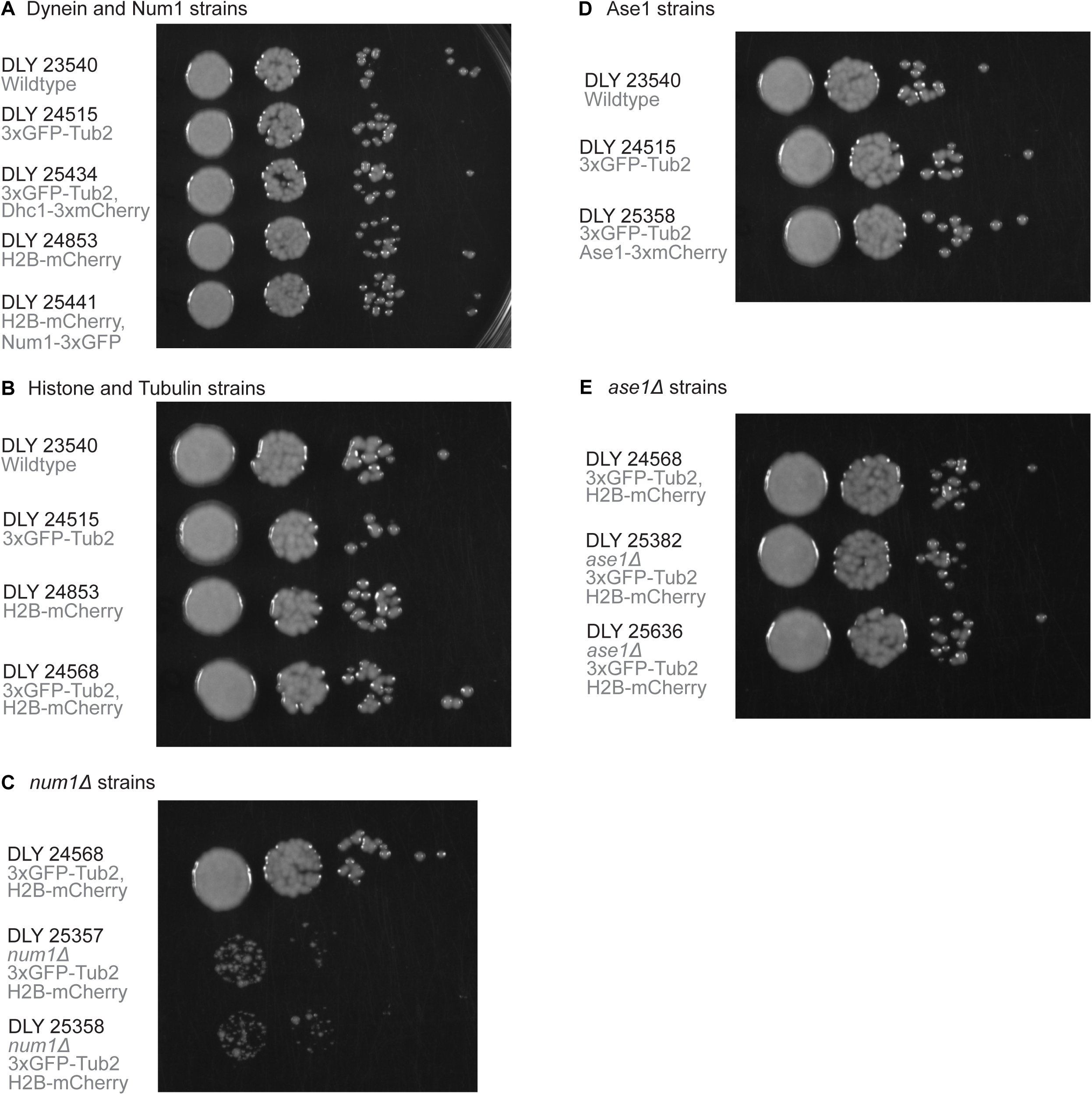
Colony growth of *A. pullulans* mutants and strains expressing fluorescent probes. Cells (5000, 500, 50, 5) of the indicated strains were spotted onto YPD and grown for two days at 25°C. *num1Δ* cells have an extreme growth defect.

**Supplemental Figure 4:**
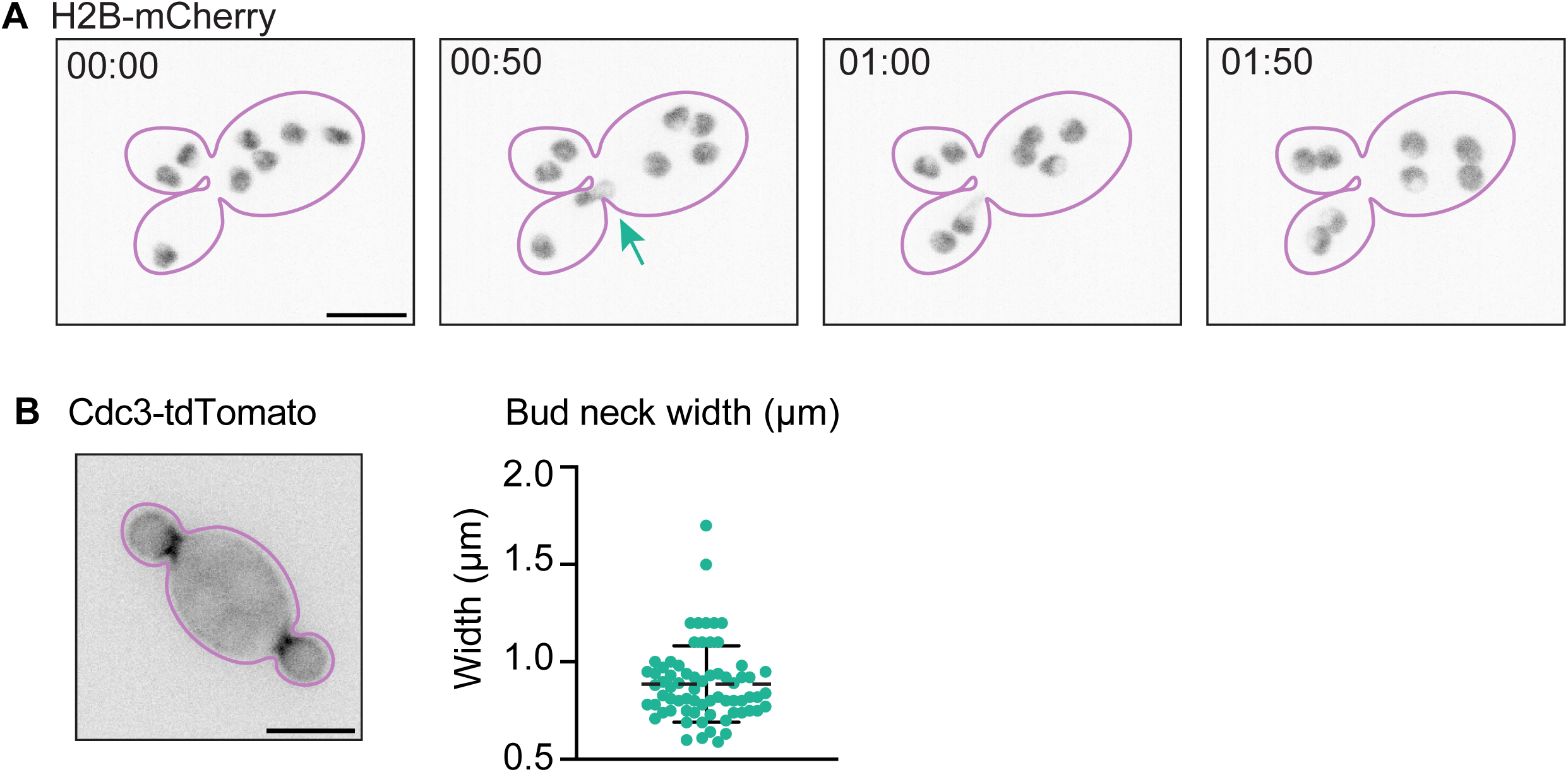
Nuclei squeeze through narrow bud necks in *A. pullulans*. (A) Inverted maximum projection time-lapse images of cells expressing H2B-mCherry (DLY24853). Nuclei appear deformed during passage through the neck (green arrow). (B) Inverted single plane image of cell expressing Cdc3-tdTomato (DLY24158). Medial plane images were used to measure the bud-neck width: each dot indicates one neck (n = 69 necks). Mean width is indicated with dashed line. Scale bars, 5 μm.

**Supplemental Table 1:**
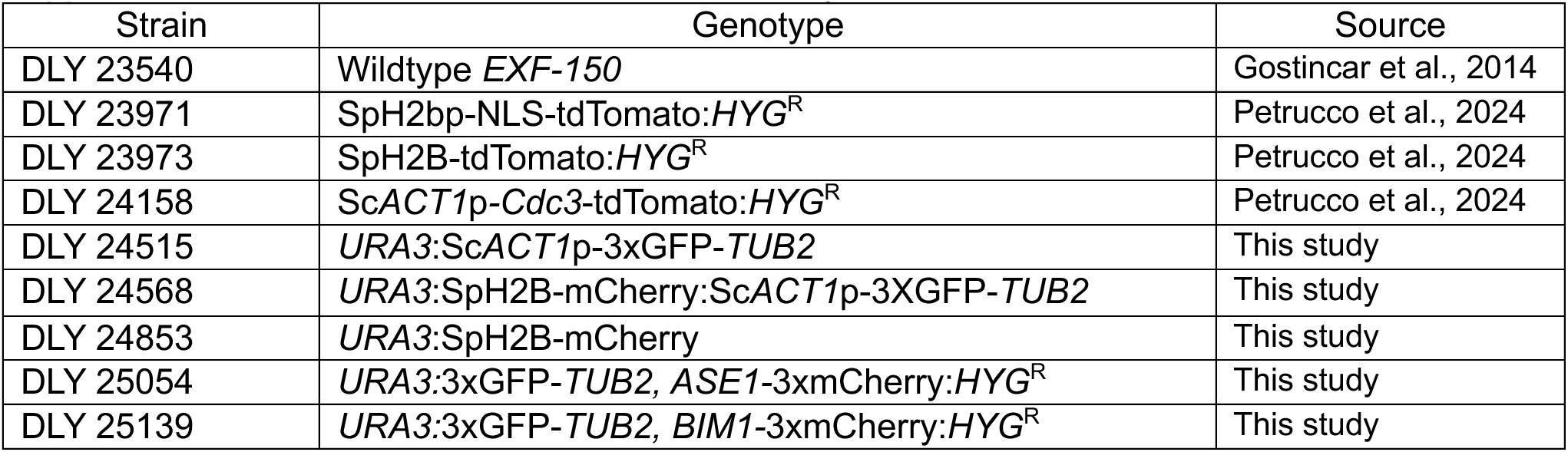
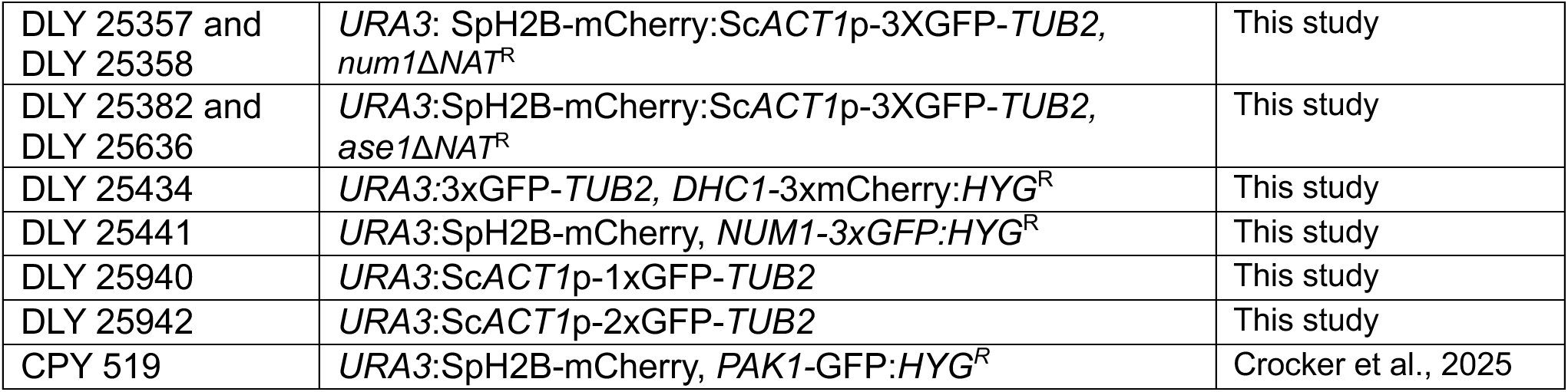
Yeast strains used in this study.

**Supplemental Table 2:**
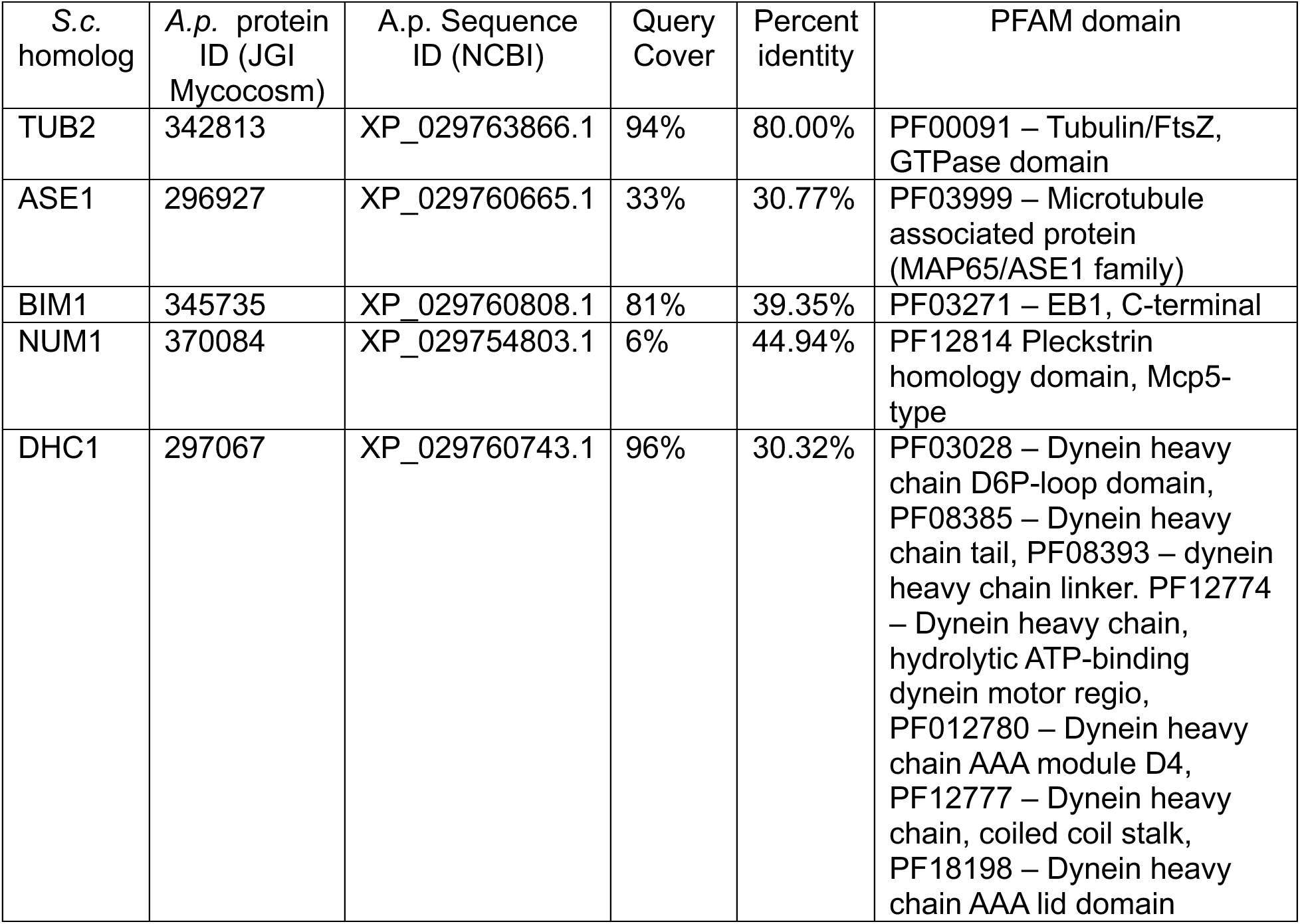
Proteins analyzed in this study.

| S.c. homolog | A.p. protein ID (JGI Mycocosm) | A.p. Sequence ID (NCBI) | Query Cover | Percent identity | PFAM domain |
| --- | --- | --- | --- | --- | --- |
| TUB2 | 342813 | XP_029763866.1 | 94% | 80.00% | PF00091 – Tubulin/FtsZ, GTPase domain |
| ASE1 | 296927 | XP_029760665.1 | 33% | 30.77% | PF03999 – Microtubule associated protein (MAP65/ASE1 family) |
| BIM1 | 345735 | XP_029760808.1 | 81% | 39.35% | PF03271 – EB1, C-terminal |
| NUM1 | 370084 | XP_029754803.1 | 6% | 44.94% | PF12814 Pleckstrin homology domain, Mcp5-type |
| DHC1 | 297067 | XP_029760743.1 | 96% | 30.32% | PF03028 – Dynein heavy chain D6P-loop domain, PF08385 – Dynein heavy chain tail, PF08393 – dynein heavy chain linker. PF12774 – Dynein heavy chain, hydrolytic ATP-binding dynein motor regio, PF012780 – Dynein heavy chain AAA module D4, PF12777 – Dynein heavy chain, coiled coil stalk, PF18198 – Dynein heavy chain AAA lid domain |

